# Experimentally-Determined Strengths of Atom-Atom (C, N, O) Interactions Responsible for Protein Self-Assembly in Water: Applications to Folding and Other Protein Processes

**DOI:** 10.1101/2020.05.26.104851

**Authors:** Xian Cheng, Irina A. Shkel, Kevin O’Connor, M. Thomas Record

## Abstract

Folding and other protein self-assembly processes are driven by favorable interactions between O, N, and C unified atoms of the polypeptide backbone and sidechains. These processes are perturbed by solutes that interact with these atoms differently than water does. C=O···HN hydrogen bonding and various π-system interactions have been better-characterized structurally or by simulations than experimentally in water, and unfavorable interactions are relatively uncharacterized. To address this situation, we previously quantified interactions of alkylureas with amide and aromatic compounds, relative to interactions with water. Analysis yielded strengths of interaction of each alkylurea with unit areas of different hybridization states of unified O, N, C atoms of amide and aromatic compounds. Here, by osmometry, we quantify interactions of ten pairs of amides selected to complete this dataset. A novel analysis yields intrinsic strengths of six favorable and four unfavorable atom-atom interactions, expressed per unit area of each atom and relative to interactions with water. The most favorable interactions are sp^2^O - sp^2^C (lone pair-π, presumably n-π*), sp^2^C - sp^2^C (π-π and/or hydrophobic), sp^2^O-sp^2^N (hydrogen bonding) and sp^3^C-sp^2^C (CH-π and/or hydrophobic). Interactions of sp^3^C with itself (hydrophobic) and with sp^2^N are modestly favorable, while sp^2^N interactions with sp^2^N and with amide/aromatic sp^2^C are modestly unfavorable. Amide sp^2^O-sp^2^O interactions and sp^2^O-sp^3^C interactions are more unfavorable, indicating the preference of amide sp^2^O to interact with water. These intrinsic interaction strengths are used to predict interactions of amides with proteins and chemical effects of amides (including urea, N-ethylpyrrolidone (NEP), and polyvinyl-pyrrolidone (PVP)) on protein stability.

**Significance:** Quantitative information about strengths of amide nitrogen-amide oxygen hydrogen bonds and π-system and hydrophobic interactions involving amide-context sp^2^ and/or sp^3^ carbons is needed to assess their contributions to specificity and stability of protein folds and assemblies in water, as well as to predict or interpret how urea and other amides interact with proteins and affect protein processes. Here we obtain this information from thermodynamic measurements of interactions between small amide molecules in water and a novel analysis that determines intrinsic strengths of atom-atom interactions, relative to water and per unit area of each atom-type present in amide compounds. These findings allow prediction or interpretation of effects of any amide on protein processes from structure, and may be useful to analyze protein interfaces.

## Introduction

Biopolymer self-assembly in water, including folding, binding, droplet formation, phase separation and formation of the functional protein and nucleic acid complexes of the cell, is driven by net-favorable interactions between C, N and O unified atoms of their biochemical functional groups, relative to interactions with water. To understand the energetics of these processes and relate thermodynamic and structural information, strengths of interaction of the different types of C, N and O unified atoms with one another, relative to their interactions with water, must be determined. Effects of biochemical solutes and noncoulombic effects of salts from the Hofmeister series on these biopolymer processes result from preferential interactions of the C, N, O atoms of the solute (and inorganic ions of the salt) with the C, N and O atoms of the biomolecule (1, 2). Quantitative information about preferential interactions of the various types of C, N and O atoms of biomolecules and solutes will therefore be useful in analyzing both self-assembly interactions and solute effects on these interactions.

Hydrogen bonding between amide sp^2^O and sp^2^N unified atoms (3-6) and the hydrophobic effect of reducing the exposure of sp^2^C and sp^3^C atoms to water (7-10), have long been recognized to be key determinants of specificity and stability of protein assemblies and complexes. In addition, *n*-π* interactions (a type of lone pair – π interaction (11)) between amide sp^2^O and amide or aromatic sp^2^C (12-17), π-π interactions of sp^2^C with sp^2^C (18, 19) and CH-π interactions of sp^3^C with sp^2^C (20-23) have been characterized by structural, spectroscopic and computational studies. Much less is known about the relative strengths of these and other amide atom-atom contacts in water, including interactions of amide sp^2^N with amide sp^2^N, sp^3^C and sp^2^C and interactions of amide sp^2^O with amide sp^2^O and sp^3^C.

Preferential interactions of biochemical solutes and Hofmeister salts with other solutes or biopolymers, relative to interactions with water, are quantified by chemical potential derivatives (∂µ_2_/∂m_3_)_T,P,m2_ = µ_23_, where the subscripts “2” and “3” refer to the two solutes, respectively, and µ_23_ = µ_32_.(1, 2). These µ_23_ values, related to transfer free energies, are determined by osmometry or solubility assays (20, 24-32). Integration of the radial distribution (33-36) of one solute in the vicinity of the other, obtained from molecular dynamics simulations (36-41), also yields µ_23_ (31).

Experimental research and analysis extending over the last decade (24-32) has shown that µ_23_ values are accurately described as a sum of contributions (*α*_3,*i*_*ASA*_*i*(2)_) from interactions of solute 3 with the accessible surface of the different types of C, N, O atoms on solute 2:

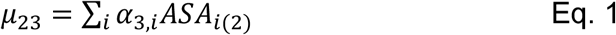

The choice of which solute to designate as component 2 or 3 is arbitrary because µ_23_ = µ_32._

In Eq. 1, each intensive quantity *α*_3,*i*_ is a thermodynamic coefficient (called a one-way alpha value) that quantifies the strength of interaction of the amide compound designated component 3 (e.g. urea) with a unit area (1 Å^2^) of one of the *i* different types (i.e. hybridization states) of C, N and O atoms on the set of amide compounds (each designated component 2), relative to interactions with water. The extensive quantity *ASA*_*i*(2)_ is the water-accessible surface area in Å^2^ of the *i* th atom type of the amide solute (component 2) whose interaction with amide solute 3 is quantified by *μ*_23_. Examples of Eq. 1 for the interaction of two amide compounds investigated here are provided in SI Eqs. S3-4.

Eq. 1 is based on the two hypotheses that contributions to µ_23_ from different weak solute-atom interactions are additive and increase in proportion to the ASA of that C, N or O atom. Additivity has been tested and validated by the analysis of sets of µ_23_ values using Eq. 1, because the size of the µ_23_ data sets greatly exceed the number of one-way alpha values determined from the analysis. ASA is found to be a better choice of extensive variable than the number of atoms or weighted number of atoms (31, 32). Using one-way alpha values *α*_3,*i*_, effects of solutes (species *3*) on biopolymer (species *2*) processes are predicted or interpreted in terms of the interaction of solute *3* with the different types of biopolymer surface area (*ASA*_*i*(2)_) exposed or buried in the process.

µ_23_ -Values can be interpreted as free energy changes for transfer of a solute from a two component solution to a three component solution in which the concentration of the other solute is 1 molal. Originally, effects of urea and osmolytes on protein stability were interpreted assuming additivity of transfer free energy contributions from the peptide backbone and each of the nineteen different amino acid side chains that are exposed to the solution in unfolding (42-46). These twenty side chain and backbone transfer free energies were obtained from amino acid and dipeptide solubility data, also assuming additivity. Analysis of urea effects on protein stability using Eq. 1 involves many fewer parameters (from as few as two (47) to four (31) or seven (26, 27) one-way alpha values (*α*_*urea,i*_), depending on the extent of coarse-graining of the ASA exposed in unfolding). One-way alpha values are interpretable in terms of the local accumulation or exclusion of the solute in the vicinity of a particular type of atom on the model compound or protein, using the solute partitioning model (SPM) (2, 20, 24-29, 48-52).

One-way alpha values are found to have fundamental chemical significance. For example, the one-way alpha value for interaction of urea with amide sp^2^O atoms (*α*_*urea,sp*2*O*_) is favorable (31) and of similar magnitude to (∼35% smaller than) that for the interaction of urea with carbonyl sp^2^O atoms of nucleobases (32). This observation indicates that hydrogen bonding interactions of the two urea sp^2^N unified atoms (hydrogen bond donors) with amide and nucleobase carbonyl sp^2^O (acceptor) must be of similar strength per unit area of sp^2^O surface and that these sp^2^N - sp^2^O interactions are more favorable than interactions of the separate atoms with water. It also indicates that these favorable sp^2^N - sp^2^O hydrogen bonding interactions contribute more to the observed one-way alpha value (*α*_*urea,sp*2*O*_) than interactions of urea sp^2^O with amide or nucleobase sp^2^O, which are expected to be quite unfavorable because these sp^2^O atoms hydrogen bond to water but have no way to interact favorably with each other.

Alkylation of urea reduces sp^2^N ASA, reducing its ability to hydrogen bond to amide and carbonyl sp^2^O, and introduces aliphatic sp^3^C which is expected to interact unfavorably with amide and carbonyl sp^2^O. Consistent with the above, as the extent of alkylation increases the one-way alpha value for interaction with amide and carbonyl sp^2^O becomes increasingly unfavorable. These trends extend to other one-way alkylurea alpha values, as discussed previously (31, 32).

The above examples lead to the hypothesis that one-way alpha values for amide compounds can be dissected into contributions from the interaction of the different types of atoms on amide solute 3 (e.g. amide sp^2^O, sp^2^N, sp^2^C for urea) with each type of atom on the set of amide solutes 2. Such a two-way dissection would quantify atom-atom interactions (designated simply as two-way alpha values) relative to water and per unit ASA of each atom.

The research reported here tests this hypothesis for the interactions of amide compounds with other amide compounds. These amide compounds display four of the most common types of unified atoms of proteins (amide sp^2^O, N, C; sp^3^C). Our previous study focused on the series of alkylated ureas, all of which have small water-accessible surface area of amide sp^2^C atoms. Here we extend the amide dataset by determining µ_23_ values for interactions of five other amides, including formamide and N-methyl formamide, which have large water-accessible surface areas (ASA) of amide sp^2^C, and malonamide and *N*-acetyl-alanine *N*-methylamide (aama), which have two amide groups and correspondingly larger ASA of amide sp^2^O.

Analysis of the combined µ_23_ dataset for amides (more than one hundred µ_23_-values) yields a set of ten two-way alpha values that quantify all possible pairwise interactions between amide sp^2^O, amide sp^2^N, sp^2^C and sp^3^C atoms of these amide compounds. We demonstrate the use of these two-way alpha values to predict µ_23_ values for interactions of any two amide compounds (with amide sp^2^O, N, C and aliphatic sp^3^C) in water from ASA information. Since proteins are of course polyamides, µ_23_ values for any amide-protein interaction (or protein-protein interaction involving only amide and hydrocarbon surface) can be predicted from these two-way alpha values, as can standard free energy derivatives 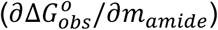 called *m*-values (equal to Δµ_23_) that quantify the effect of any amide solute on any protein process in which the change in ASA is primarily from amide and hydrocarbon atoms (as is the case for protein folding). In addition, the rank order of two-way alpha values and relative strengths of these atom-atom preferential interactions should provide a useful starting point for assessing the contributions of different atom-atom contacts to the stability of a protein interface.

## Results

### Analysis to Determine Atom-atom Interactions from Solute-Atom Interactions

For interactions of a series of urea and alkyl ureas (component 3) with amide compounds (component 2), analyzed by Eq. 1 as summarized above, each of the solute (3) - atom (i) one-way alpha values (***α***_**3,*i***_) exhibited a regular progression with increasing alkylation (and reduced exposure of amide nitrogen) of the urea (Fig. S1; (31)). Motivated by this observation, here we test the hypothesis that each of these one-way alpha values can be dissected into additive, ASA-based contributions from the interaction of the different types of atoms on amide solute 3 (sp^2^O, N, C and sp^3^C) with the i-th type of atom (also sp^2^O, N, C or sp^3^C) on amide solute 2.

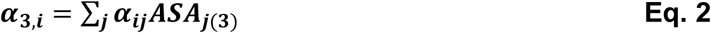

Eq. 2 for the one-way alpha value *α*_3,*i*_ is completely analogous to Eq. 1 for µ_23_. In Eq. 2, each intensive quantity ***α***_***ij***_ is the strength of interaction of a unit area of atom j of solute 3 with a unit area of atom i of solute 2, and the corresponding extensive quantity *ASA*_*j*(3)_ is the accessible surface area of atom type j on solute 3. For the amide-amide interactions of interest here, an example with all the individual terms in the sum in Eq. 2 is provided in SI Eqs. S5-6. The hypotheses of additivity and ASA-dependence of the contributions *α*_*ij*_ *ASA*_*j*(3)_ to the one-way alpha value *α*_3,*i*_ are tested concurrently with the determination of two-way alpha values (*α*_*ij*_) because the number of equations (like Eq. 2) greatly exceeds the number of unknowns (*α*_*ij*_) being determined (see below).

A straightforward test of Eq. 2 is provided by the comparison of one-way alpha-values *α*_3,*i*_ for selected pairs of amide solutes that differ primarily in ASA of one atom type (j). For these situations, a semiquantitative estimate of the two way alpha value *α*_*ij*_ is obtained from the difference in one-way alpha values Δ*α*_3,*i*_ and the ASA difference for atom type j (Δ*ASA*_*j*(3)_)

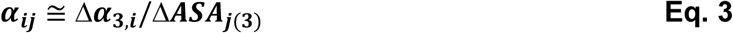

An example of this calculation is provided in SI (Eqs. S7-10) to clarify the notation.

Combination of Eqs. 1 and 2 gives the proposed dissection of any solute-model compound µ_23_ – value into contributions from the interactions of accessible atoms of the solute with accessible atoms of the model compound:

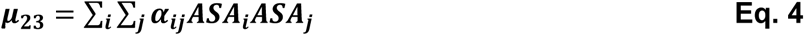

where ***α***_***ij***_= ***α***_***ji***_. The complete set of terms in this double sum for the amide compounds investigated here is given in SI as Eq. S11.

As an interpretation of one term in Eq. 4, consider the contribution to µ_23_ from the interaction of amide sp^2^N of one amide solute (component 2) with amide sp^2^O on a second amide solute (component 3), relative to interactions with water, given by ***α***_***sp*2*N*−*sp*2*O***_***ASA***_**2,*sp*2*N***_***ASA***_**3,*sp*2*O***_. The product of ASA values is proportional to the probability that a contact between the two solutes involves sp^2^N atom(s) of solute component 2 and sp^2^O atom(s) of solute component 3, and the two-way alpha value ***α***_***sp*2*N*−*sp*2*O***_ is the strength of that interaction per unit ASA of both atom types, again relative to water. These two-way alpha values are useful to predict or interpret µ_23_ values for interactions of amides for which one-way alpha values are not available, and to predict or interpret effects of these amides on protein processes in terms of structural information. Two-way alpha values may also be useful in analyses of atom-atom interactions in protein assemblies and in binding interfaces.

### VPO Determinations of Interactions of Amides with Large ASA of Amide sp^2^C and sp^2^O

Previously we determined one-way alpha values *α*_3,*i*_ (Eq 1) quantifying interactions of urea and six alkyl urea solutes with the types of unified atoms of amide (sp^2^O, sp^2^N, sp^2^C; sp^3^C) and aromatic hydrocarbon (sp^2^C) compounds using osmometric and solubility studies (31). Trends in these one-way alpha values with increasing alkylation of the urea showed which atom-atom interactions are favorable and which are unfavorable. However, preliminary tests of Eqs. 2 and 4 using the ninety-five *μ*_23_ values from this previous study revealed that these were insufficient to accurately quantify all atom-atom interactions (two-way alpha values) involving amide sp^2^O and/or sp^2^C.

Here we determine *μ*_23_ values by osmometry for an additional ten interactions of five amide compounds, including interactions of two amides with large ASA of amide sp^2^C (formamide, N-methylformamide) with one another and with two amides with large ASA of amide sp^2^O (malonamide, aama). Interactions of these four amides with propionamide are also determined. In addition to their significance for the two-way analysis proposed here, these measurements also permit the determination of one-way alpha values for the interactions of these five amides with amide sp^2^O, sp^2^N, sp^2^C and aliphatic sp^3^C atoms, increasing the number of amide compounds for which one-way alpha values (Eq 1) are available from seven to twelve.

For uncharged solutes at concentrations up to ∼1 molal, the difference ΔOsm = Osm(m_2_,m_3_) – Osm(m_2_) – Osm(m_3_) between the osmolality (Osm) of a three component solution and the two corresponding two-component solutions is proportional to the product of solute molal concentrations (m_2_m_3_) with proportionality constant μ_23_/RT (31).

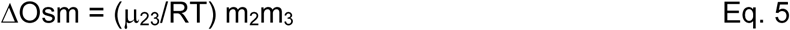

If ΔOsm is negative, μ_23_ is negative, μ_2_ decreases with increasing m_3_, and the interaction of the two solutes is favorable.

For each of the ten pairs of amides investigated here, ΔOsm is plotted vs. m_2_m_3_ in the panels of Fig. 1. All µ_23_ values are negative, indicating favorable interactions between all ten pairs of amides investigated here. Values of µ_23_ at 23 °C obtained from these slopes are listed in Table S1. Of these, the interaction of malonamide and aama (middle panel of Fig. 1) is the least favorable(µ_23_ = - 8.6 ± 2.2 cal mol^-1^ molal^-1^) and the interaction of propionamide and N-methyl formamide (top panel of Fig. 1) is the most favorable (µ_23_ = - 102 ± 1.9 cal mol^-1^ molal^-1^). Though there is substantial scatter in the data for some pairs of amides, slopes µ_23_/RT are quite well determined (see Table S1) because the intercept is constrained to be zero.

**Figure 1.**
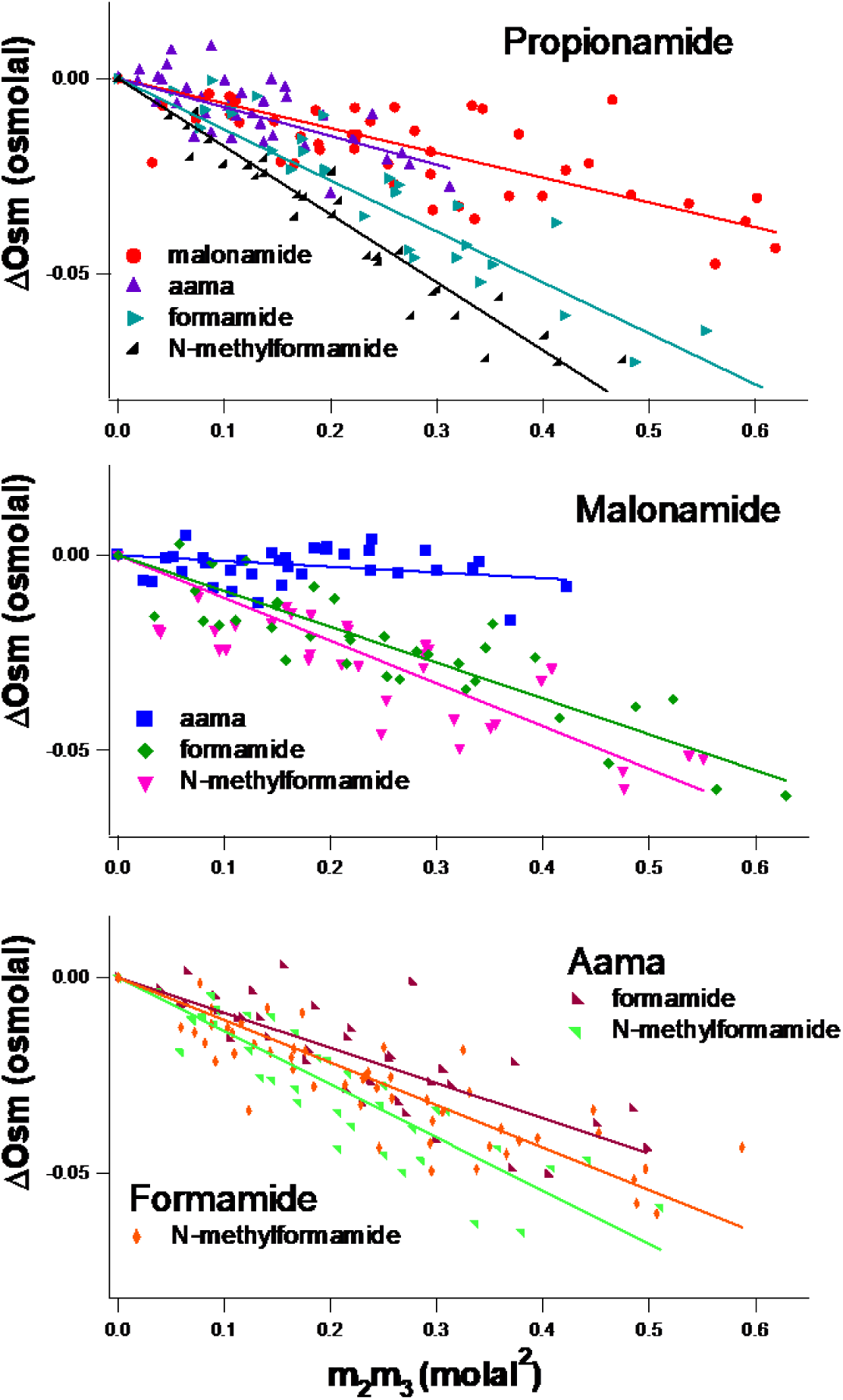
Osmometric Determinations of Preferential Interactions of Pairs of Amide Compounds in Water. Osmolality differences ΔOsm = Osm(m_2_,m_3_) – (Osm(m_2_) + Osm(m_3_)) between a three-component solution of two amide compounds and the two corresponding two-component solutions, determined by VPO at 23 °C, are plotted vs. the product of molal concentrations (m_2_m_3_) of the two amides (Eq. 5). Slopes of linear fits with zero intercept yield chemical potential derivatives 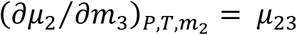 quantifying preferential interactions between the two amides. aama: N-Acetylalanine N-methylamide.

### One-way Alpha values for Interactions of Five Amide Solutes with Amide O, N and C Atoms

Analysis of the ten µ_23_-values determined from Fig. 1 together with previous results for the interactions of these five amides with other amides (31) by Eq. 1 yields one-way alpha values *α*_3,*i*_ for interactions of these five amides with each of the four types of unified atoms of amide compounds. These one-way alpha values are plotted as bar graphs in Fig. 2 and listed in Table S2. Fig S1 compares one-way alpha value one-way alpha values for all 12 amide compounds investigated to date. Figs. 2 and S1 show that all amide compounds investigated here and previously interact favorably with amide sp^2^C, amide sp^2^N, and aliphatic sp^3^C, and that all but urea and formamide interact unfavorably with amide sp^2^O.

**Figure 2.**
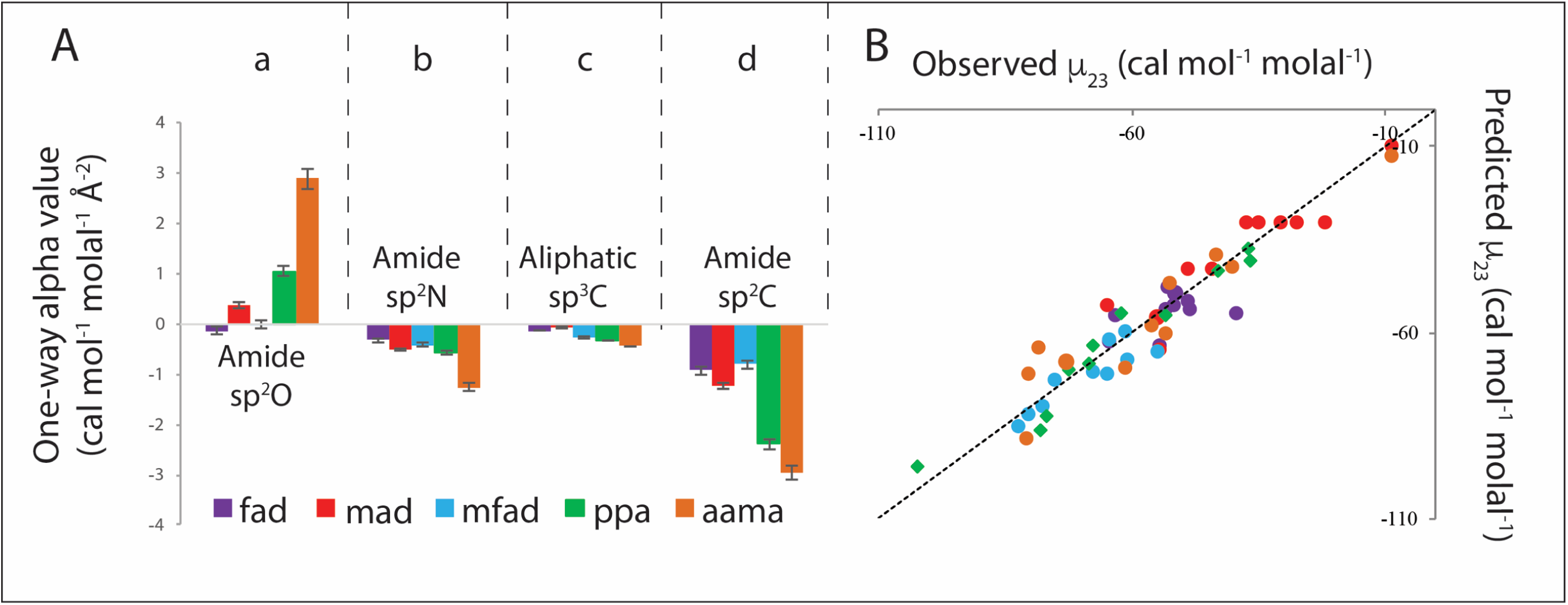
**A**: **Strengths of Interaction of Five Amides (formamide (fad), N-methylformamide (mfad), malonamide (mad), propionamide (ppa) and Acetyl-L-ala-methylamide (aama)) with Amide and Hydrocarbon Unified Atoms.** Bar graphs compare interaction potentials (α-values; Table S2)) quantifying interactions of these five amide compounds with a unit area of amide sp^2^O, amide sp^2^N, aliphatic sp^3^C, and amide sp^2^C at 23°C. Favorable interactions have negative one-way alpha values while unfavorable interactions have positive one-way alpha values. **B**: Comparison of Predicted and Observed µ_23_ Values for Pairwise Interactions of these Five Amide Compounds at 23 °C. Predictions of µ_23_ use one-way alpha values for these five amide compounds with amide sp^2^O, amide sp^2^N, amide sp^2^C and aliphatic sp^3^ C from Panel A and Table S2. Color scheme is that of Panel A. Observed µ_23_ values are from Tables S1 and S3. The line represents equality of predicted and observed values.

### Strengths of Pairwise Interactions of Amide sp^2^O, N, C and Aliphatic sp^3^C Unified Atoms

All one hundred and five µ_23_ values (Tables S3, S4) for interactions of twelve different amide compounds with each other, and in some cases with naphthalene and/or anthracene, were analyzed using Eq. 4 to obtain ten two-way alpha values (α_sp2O,sp2O_, α_sp2O,sp2N_, α_sp2O, sp2C_, α_sp2O,sp3C_; α_sp2N,sp2N_, α_sp2N,sp2C_, α_sp2N,sp3C_; α_sp2C,sp2C_, α_sp2C,sp3C_; α_sp3C,sp3C_). The previous one-way (31) analysis revealed that interactions of sp^2^C atoms of aromatic hydrocarbons and of amides are similar if not identical, and they are analyzed together here. (Alternative analyses of sp^2^C presented in SI and discussed below justify this treatment.) In this analysis of all amide-amide and amide-aromatic µ_23_ values, the number of equations (105 applications of Eq. 4) exceeds the number of unknowns (10 two-way alpha values) by more than ten-fold, making them highly overdetermined. The additivity and ASA-dependence of contributions to µ_23_ underlying Eq. 4 are tested quantitatively by comparison of predicted and observed µ_23_ values (Tables S3, S4, Fig. 3B), and semi-quantitatively from differences in one-way alpha values for amides differing primarily in ASA of one type of unified atom (see below).

**Figure 3.**
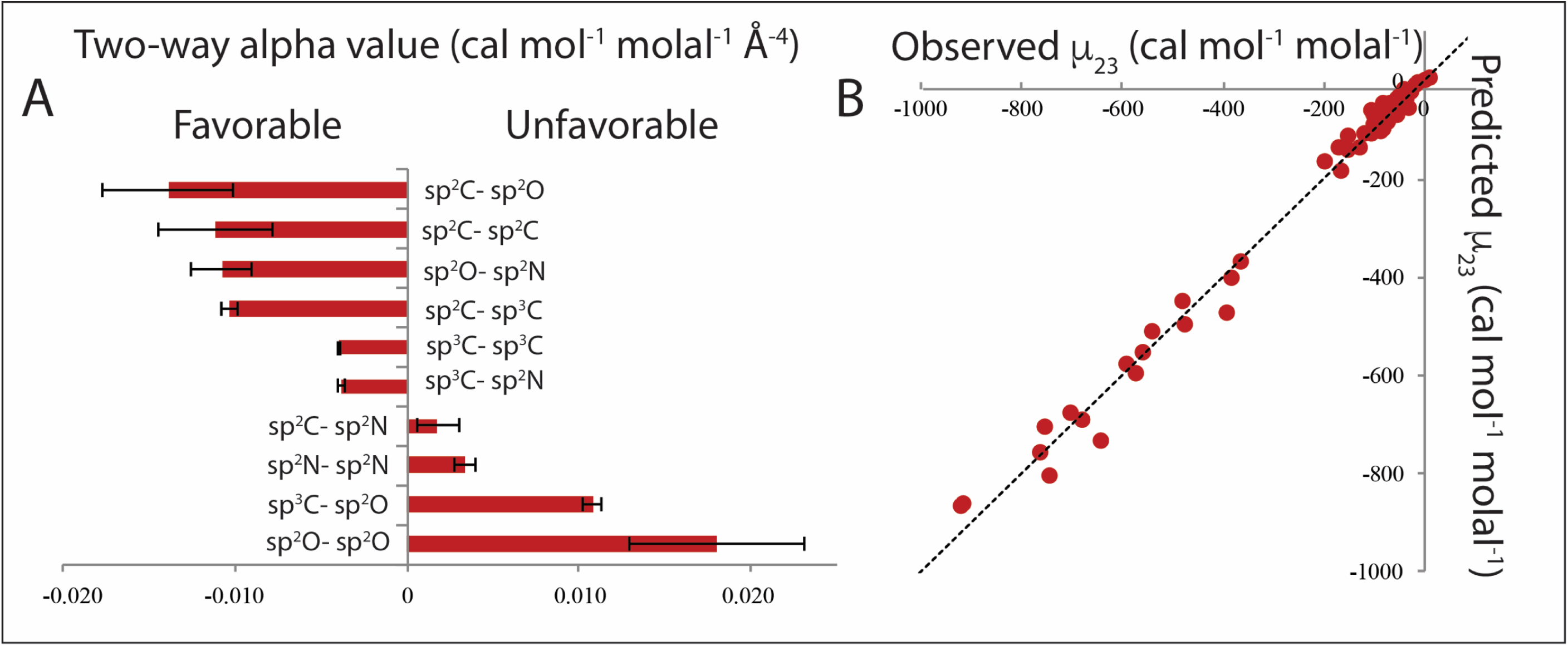
Amide Compound Atom-Atom Interaction Strengths and their Ability to Predict µ_23_ Values. A) Two-way alpha values (Table 1) quantifying interactions of pairs of amide atoms shown, relative to interactions with water at 23 °C. sp^2^C is for combined amide and aromatic sp^2^C. Negative alpha values indicate favorable interactions. B) Comparison of predicted and observed µ_23_ values for interactions of each pair of amide compounds investigated at 23 °C. Predictions of µ_23_ use two-way alpha values in Table 1. Observed µ_23_ values are from Tables S1 and S3. The line represents equality of predicted and observed values.

Results of this analysis (two-way alpha values) quantifying the pairwise interactions of amide sp^2^O, sp^2^N, sp^2^C and aliphatic sp^3^C unified atoms with one another are listed in Table 1. These ten two-way alpha values are also plotted in the bar graph of Fig. 3A in ranked order from the most negative (most favorable interactions relative to interactions with water) to the most positive (most unfavorable interactions). Uncertainties in these two-way alpha values range from 3% to 30% except for the small-magnitude interaction of sp^2^C with sp^2^N, where the uncertainty is larger (∼70%). These uncertainties do not affect the semi-quantitative conclusions of this research.

**Table 1.** Ranked Intrinsic Strengths of Atom-Atom Interactions of Amide Compounds.

| Interaction Type | Amide Atom |  | Strength<sup>a</sup> |  |
| --- | --- | --- | --- | --- |
|  | i | j | Two-way alpha value<br>(cal mol<sup>-1</sup> molal<sup>-1</sup> Å<sup>-4</sup>) | Relative to<br>sp<sup>3</sup>C – sp<sup>3</sup>C |
| **Aliphatic sp<sup>3</sup>C Interactions** |  |  |  |  |
|  | sp<sup>3</sup>C | sp<sup>2</sup>C | -0.0103 ± 0.0005 | 2.6 |
|  | sp<sup>3</sup>C | sp<sup>3</sup>C | -0.0039 ± 0.0001 | 1.0 |
|  | sp<sup>3</sup>C | sp<sup>2</sup>N | -0.0038 ± 0.0002 | 1.0 |
|  | sp<sup>3</sup>C | sp<sup>2</sup>O | 0.0108 ± 0.0005 | (2.8)<sup>b</sup> |
| **Amide sp<sup>2</sup>C Interactions** |  |  |  |  |
|  | sp<sup>2</sup>C | sp<sup>2</sup>O | -0.0139 ± 0.0038 | 3.6 |
|  | sp<sup>2</sup>C | sp<sup>2</sup>C | -0.0111 ± 0.0033 | 2.9 |
|  | sp<sup>2</sup>C | sp<sup>3</sup>C | -0.0103 ± 0.0005 | 2.6 |
|  | sp<sup>2</sup>C | sp<sup>2</sup>N | 0.0018 ± 0.0013 | (0.5)<sup>b</sup> |
| **Amide sp<sup>2</sup>O Interactions** |  |  |  |  |
|  | sp<sup>2</sup>O | sp<sup>2</sup>C | -0.0139 ± 0.0038 | 3.6 |
|  | sp<sup>2</sup>O | sp<sup>2</sup>N | -0.0108 ± 0.0017 | 2.8 |
|  | sp<sup>2</sup>O | sp<sup>3</sup>C | 0.0108 ± 0.0005 | (2.8)<sup>b</sup> |
|  | sp<sup>2</sup>O | sp<sup>2</sup>O | 0.0181 ± 0.0051 | (4.6)<sup>b</sup> |
| **Amide sp<sup>2</sup>N Interactions** |  |  |  |  |
|  | sp<sup>2</sup>N | sp<sup>2</sup>O | -0.0108 ± 0.0017 | 2.8 |
|  | sp<sup>2</sup>N | sp<sup>3</sup>C | -0.0038 ± 0.0002 | 1.0 |
|  | sp<sup>2</sup>N | sp<sup>2</sup>C | 0.0018 ± 0.0013 | (0.5)<sup>b</sup> |
|  | sp<sup>2</sup>N | sp<sup>2</sup>N | 0.0034 ± 0.0006 | (0.9)<sup>b</sup> |

<sup>a</sup> Interactions are symmetric ( $\alpha_{ij} = \alpha_{ji}$ ) so only 10 of the 16 two-way alpha values listed are unique.
<sup>b</sup> Unfavorable interaction strengths (indicated by parentheses) are expressed relative to the magnitude of the sp<sup>3</sup>C – sp<sup>3</sup>C interaction.

Six of the ten atom-atom interactions in Table 1 are favorable, with negative two-way alpha values. The four most favorable interactions, of similar strength when expressed per unit area of each unified atom, are sp^2^O - sp^2^C, sp^2^C - sp^2^C, sp^2^O - sp^2^N and sp^2^C - sp^3^C. Two-way alpha values for these four interactions are about three times more negative than two-way alpha values for sp^3^C - sp^3^C and sp^3^C - sp^2^N interactions. The most unfavorable interaction in this set is sp^2^O - sp^2^O. Sp^2^O - sp^3^C and sp^2^N - sp^2^N interactions are modestly unfavorable, while the sp^2^N - sp^2^C interaction is slightly unfavorable. The signs and relative magnitudes of these interaction strengths, relative to interactions with water, make good chemical sense as discussed below and in SI. We therefore conclude that these two-way alpha values not only are useful to predict and interpret interactions of amide solutes and amide effects on protein processes, but also have fundamental chemical significance for understanding the weak interactions that drive self-assembly.

### Comparison of Two-way Alpha Values Obtained by Fitting with Estimates from One-way Alpha Values for Amide Pairs Differing Primarily in ASA of One Atom Type

Ethylurea and 1,1-diethylurea differ from methylurea and 1,1-dimethylurea primarily in aliphatic sp^3^C ASA. The sp^3^C ASA difference between the dialkylated ureas (64 Å^2^) is about twice as large for the monoalkylated ureas (36 Å^2^). These sp^3^C ASA differences are 85-90% of the total magnitude of ASA differences for these pairs of compounds. Applying Eq. 3 to estimate two-way alpha values for the interactions of aliphatic sp^3^C atoms with amide sp^2^O, N and C atoms from differences in the corresponding one-way alpha values for these pairs of amide compounds and these differences in sp^3^C ASA yields the direct estimates in Table S5. For the dialkyl ureas, where the differences in one-way alpha values are larger and better determined, estimates of two-way alpha values for interactions of sp^3^C with all four atom types agree quantitatively with those obtained from global fitting, differing by less than the propagated uncertainty in the fitted values. For the monoalkyl ureas, agreement is semiquantitative, with deviations of 25-40%.

1,3-Diethylurea differs from proprionamide primarily in amide sp^2^N ASA (20 Å^2^, which is 84% of the total magnitude of ASA differences for this pair of compounds). Table S5 shows a range of agreement between estimates of two-way alpha values from Eq. 3 and best fit two-way alpha values. Near-quantitative agreement of two-way alpha values is obtained for interactions of amide sp^2^N with amide sp^2^O and aliphatic sp^3^C, while estimates for interactions with amide sp^2^N and amide sp^2^C agree with best-fit two-way alpha values in sign but not in magnitude. With the exception of the strongly favorable amide sp^2^N - amide sp^2^O interaction, atom-atom interactions involving amide sp^2^N are weak, which affects the ability to determine them by this difference approach. Finally, the diamide Acetyl-L-ala-methylamide (aama) differs from 1,3-diethylurea (1,3-deu) primarily in amide sp^2^O ASA (34 Å^2^, which is 63% of the total magnitude of ASA differences for this pair of compounds). Estimates of two-way alpha values for interactions of sp^2^O from differences in one-way alpha values (Eq. 3), listed in Table S5, agree with best-fit two-way alpha values within 10-30%. This level of agreement is obtained because interactions involving sp^2^O are the strongest of any atom type, and hence are dominant even for this situation where the ASA difference is only 63% sp^2^O.

### Comparison of Observed µ_23_ Values with Predictions from Alpha Values and ASA

Fig. 3B compares observed µ_23_ values for interactions of the series of urea and amide solutes with each other with those predicted from two-way alpha values (Tables 1) and ASA information (31) using Eq.3. All observed and predicted µ_23_ values are listed in Tables S3 and S4. For amide-amide interactions, predicted and observed µ_23_ values are in good agreement within the combined uncertainties (± 1 SD, typically 15%) for about 90% of the interactions investigated (94 out of 105).

Table S3 also lists predicted µ_23_ values for the amide compounds in the dataset calculated from one-way alpha values of five amide solutes (formamide, N-methylformamide, malonamide, propionamide and Acetyl-L-ala-methylamide (aama)) with the four amide atoms (Table S2) and ASA information using Eq. 2. Comparison in Table S4 of predicted µ_23_ values for the interaction of two amides using two-way alpha values (Table 1) with predictions using the sets of one-way alpha values determined for the two amides (Table S3) reveals that agreement with experimental data is equally good for both sets of alpha values. Comparison in Figure S2 (one-way alpha values are calculated based on combination of amide and aromatic sp^2^C) and Table S6 of one-way alpha values which are calculated from µ_23_ values with predictions of one-way alpha values from two-way alpha values shows a good agreement with each other.

## Discussion

### Chemical Significance of Amide Two-way Alpha Values

#### Interactions of Aliphatic sp^3^C with Aliphatic sp^3^C, Amide/Aromatic sp^2^C and Amide sp^2^N, O

Interactions of aliphatic sp^3^C atoms of amide compounds with aliphatic sp^3^C, amide sp^2^C and amide sp^2^N atoms of other amides are favorable, while interactions with amide sp^2^O are unfavorable, relative to interactions with water. Strengths (two-way alpha values) of these preferential interactions, expressed per unit area of each atom type (Table 1), span a wide range. These two-way alpha values are well-determined from the global fitting, with uncertainties of 3-5%.

The preferential interaction of sp^3^C with sp^2^C, quantified per unit ASA of each atom type, is one of the four most favorable atom-atom interactions characterized here. This is often called a CH-π interaction, and should also involve a hydrophobic effect from the burial of sp^3^C and sp^2^C ASA when it occurs. From Table 1 two-way alpha values, the strength of a favorable sp^3^C - sp^2^C interaction is almost three times that of a sp^3^C-sp^3^C interaction, which is presumably driven by a hydrophobic effect from removing sp^3^C ASA from water. Interpreted most simply, this comparison indicates that the CH-π component of the favorable interaction of aliphatic sp^3^C with amide or aromatic sp^2^C contributes more than the hydrophobic component of this interaction.

Two-way alpha values in Table 1 also reveal that the sp^3^C - sp^2^N preferential interaction is about as favorable as the sp^3^C - sp^3^C interaction. Because sp^2^N unified atoms are expected to interact more favorably with water than sp^3^C atoms do, it follows that the intrinsic interaction of sp^3^C with sp^2^N (i.e. not relative to water) is more favorable than the intrinsic interaction of sp^3^C with sp^3^C.

The sp^3^C - sp^2^O interaction in water is highly unfavorable, with an α -value that is equal in magnitude and opposite in sign to the sp^3^C - sp^2^C interaction. An unfavorable interaction means that intrinsic interactions of the unified sp^3^C and sp^2^O atoms with water are more favorable than the intrinsic sp^3^C - sp^2^O interaction. The sp^3^C - sp^2^O interaction is unfavorable because the intrinsic interaction of water with sp^2^O is favorable while the intrinsic interaction of sp^3^C with amide sp^2^O is probably comparably unfavorable to its intrinsic interaction with water.

#### Amide/aromatic sp^2^C Interactions

From Table 1, interactions of amide/aromatic sp^2^C atoms with amide sp^2^O, aliphatic sp^2^C, and amide sp^3^C atoms of other amides are all very favorable, while interactions with amide sp^2^N are slightly unfavorable, relative to interactions with water. Overall, amide/aromatic sp^2^C atoms interact more favorably with the atoms of amide compounds than any other atom type in Table 1. These sp^2^C two-way alpha values are not as accurately known as sp^3^C two-way alpha values. Except for the sp^2^C - sp^3^C interaction (5% uncertainty), uncertainties are 27% for interactions with sp^2^O and sp^2^C and 70% for the very weak interaction with sp^2^N.

The sp^2^C-sp^2^O interaction is the most favorable interaction quantified here, with a two-way alpha values which is about 3 times as favorable as for hydrophobic sp^3^C - sp^3^C, which we take as a reference. In all likelihood the sp^2^C-sp^2^O interaction is a n – π* interaction (one example of a lone pair (lp) – π interaction (11)) involving n-shell electrons of amide sp^2^O and the π system of the amide group or aromatic ring, as characterized previously in structural and spectroscopic studies and MD simulations (12-17). The observation that a single two-way alpha value quantifies this interaction for both amide and aromatic sp^2^C is a compelling argument for the use of ASA in this analysis. This two-way alpha value is very similar to that deduced from the one-way alpha value for the interaction of naphthalene with amide sp^2^O ((31); see SI Table S7). Water forms hydrogen bonds to amide sp^2^O atoms and presumably participates in a lone pair – π interaction with sp^2^C atoms, so the strength of the sp^2^O - sp^2^C interaction in Table 1 is relative to these competitive interactions involving water.

Comparison of two-way alpha values in Table 1 reveals that the sp^2^C - sp^2^C interaction is about as favorable, per unit area of each participant, as the sp^2^C - sp^3^C interaction discussed above. Therefore it is likely that the π – π component of the sp^2^C - sp^2^C interaction, expressed per unit area of each participant, is similar in strength to the CH-π interaction and contributes about twice as much as the hydrophobic effect to the favorable sp^2^C - sp^2^C interaction.

The un-named interaction of amide/aromatic sp^2^C with amide sp^2^N is very marginally unfavorable. This interaction is not as favorable as the sp^3^C - sp^2^N interaction, probably because the intrinsic interaction of water oxygen lone pairs with the sp^2^C π system is more favorable than the interaction of water with sp^3^C. Even so, because it is only marginally unfavorable, there should be no significant free energy penalty for forming contacts between amide/aromatic sp^2^C and amide sp^2^N in a protein interface.

#### Amide *sp*^*2*^*O Interactions*

From Table 1, interactions of amide sp^2^O atoms with amide and aromatic sp^2^C and amide sp^2^N atoms are both very favorable while amide sp^2^O interactions with aliphatic sp^3^C and amide sp^3^O are very unfavorable, relative to interactions with water. Uncertainties in these two-way alpha values are moderate, ranging from 5% for sp^2^O - sp^3^C to 28% for sp^2^O - sp^2^C and sp^2^O - sp^2^O.

The favorable interaction of amide sp^2^O with amide-aromatic sp^2^C is discussed above. The similarly favorable interaction of amide sp^2^O with amide sp^2^N in water is almost certainly the NH—O=C hydrogen bond interaction in which the unified amide sp^2^N atom is the donor and the sp^2^O atom is the acceptor (3-6). The amide sp^2^O -amide sp^2^O interaction is almost twice as unfavorable as the amide sp^2^O – aliphatic sp^3^C interaction because of the very favourable intrinsic interaction of amide sp^2^O with water, which contributes twice as much in magnitude to the two-way alpha value for sp^2^O - sp^2^O as for sp^2^O - sp^3^C.

#### Amide *sp*^*2*^*N Interactions*

From Table 1, the amide sp^2^N-amide sp^2^O interaction is the most favorable interaction involving amide sp^2^N, while the amide sp^2^N-aliphatic sp^3^C interaction is modestly favorable, and the amide sp^2^N interactions with amide-aromatic sp^2^C and amide sp^2^N is slightly unfavorable, relative to interactions with water. Uncertainties in two-way alpha values for interactions involving amide sp^2^N are small (5% to 18%) except for the very weak interaction with sp^2^C. All these interactions except amide sp^2^N-amide sp^2^N are discussed above.

The amide sp^2^N - amide sp^2^N interaction, which very likely is the NH···N hydrogen bond, is modestly unfavorable, indicating that hydrogen bonding of amide sp^2^N with water is intrinsically more favorable. Consistent with this, NH···N hydrogen bonds are seldom observed in protein secondary structures, except involving proline (53). However, a hydrogen bond between unified N atoms of heterocyclic aromatic rings occurs in both AT (also AU) and GC base pairs of nucleic acid duplexes.

#### *Using Two-Way* Alpha Values to *Predict Amide, Polyamide Effects on Biopolymer Processes*

##### a) *Predicting m-Values* for *Urea and other Amide Solutes*

One of the significant applications of two-way alpha values for amide compound atom-atom interactions is to predict or interpret effects (*m*-values) of urea or any other amide solute on protein processes (26, 28) in terms of ASA information for the amide solute and ΔASA information for each type of unified protein atom using Eqs. 2 and 5. Recently, we analyzed urea *m*-values for unfolding of globular proteins using urea one-way alpha values for amide sp^2^O, amide sp^2^N, aliphatic sp^3^C and aromatic sp^2^C and ΔASA information assuming an extended chain model of the unfolded state. Generally good agreement is obtained between experimental *m*-values and *m*-values predicted either using these four major protein atom types or using all seven protein atom types (26). Figure S3 shows that use of two-way alpha values from Table 1 yields predicted *m*-values which agree well with the two one-way predictions and with experimental *m*-values.

##### b) Predicting Chemical Contributions to Interactions of the Polyamide PVP with Protein Surfaces and Effects of PVP on Protein Processes; Comparison with PEG

The water-soluble, flexibly-coiling polyamide polyvinylpyrrolidine (PVP), available in several different molecular weight ranges, has occasionally been used in place of the polyether PEG (polyethylene glycol) as a “macromolecular crowder” in studies of protein stability and interactions under conditions of high volume occupancy. The two-way alpha values from Table 1 are useful to predict the chemical interaction of PVP and its model monomer (N-ethyl pyrrolidone, NEP) with proteins and compare PVP with PEG. For PEG, where a wide range of molecular weights from monomer to oligomers and polymers is available, we previously determined the chemical interactions of end and interior groups of PEG with the different types of protein atoms (29) and separated chemical (preferential interaction) and physical (excluded volume) effects of PEG oligomers and polymers on protein (54) and nucleic acid (55) processes. Short oligomers of PVP are not commercially available to determine preferential interactions with protein atoms as done for PEG, but since NEP and PVP are amides, their chemical interactions with protein atoms and their chemical effects on protein processes can be predicted from the two-way alpha values obtained here and reported in Table 1.

For polymeric PVP with an average degree of polymerization *N*_3_ greater than about 20 residues, the per-residue interaction with another solute (component 2), i.e. 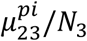, is well approximated as the interaction of an interior PVP residue, neglecting differences between end and interior residues:

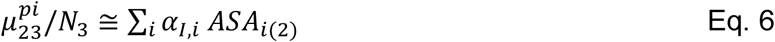

The corresponding expression for PVP oligomers where contributions from the end groups should be treated separately is given in SI. In Eq. 6, *α*_*I,i*_ quantifies the strength of interaction of an interior PVP residue with the i-th type of atom on solute 2, with accessible surface area *ASA*_*i*(2)_. While neither these *α*_*I,i*_ values for PVP nor the corresponding *α*_*NEP,i*_ values for NEP have been determined directly, both are readily predicted using Eq. 2 from the two-way alpha values in Table 1 and ASA information for PVP interior residues and NEP. Table S8 summarizes this ASA information, and Table S9 lists predicted one-way alpha values (*α*_*I,i*_, *α*_*NEP,i*_ and the PVP end-group one-way alpha value *α*_*E,i*_) and compares these PVP residue one-way alpha values with those obtained previously (29) for PEG residues.

Table S9 shows a similar pattern of interactions of PVP and PEG interior residues with the most common types of protein atoms. Chemical interactions of both PVP and PEG residues with protein aliphatic sp^3^C, amide sp^2^N and amide sp^2^C are favorable, while interactions with protein amide sp^2^O are unfavorable. Interactions of PVP interior residues with aliphatic sp^3^C and amide sp^2^O are about 2-fold stronger than for PEG, while PVP interactions with protein amide sp^2^C and amide sp^2^ N are about 1.6- and 1.2-fold stronger than for PEG.

Table S10 predicts that PVP monomer (NEP) destabilizes globular proteins, in agreement with its observed destabilization of CI2 (56). NEP is predicted to stabilize α-helices; this difference results from the very different composition of the ASA exposed in unfolding α-helices (predominantly amide, weighted toward amide sp^2^O) as compared to unfolding globular proteins (predominantly sp^3^C). By contrast, PEG monomer (ethylene glycol) and dimer (diethylene glycol) are predicted and observed to stabilize globular proteins, while tri- and tetraethylene glycol are destabilizing (29). Polymeric PVP is observed to stabilize CI2 (57, 58), which we conclude is because the predicted stabilizing excluded volume effect (54) counterbalances the predicted small destabilizing chemical effect (SI text and Table S10).

Hence PVP exerts chemical effects which differ only modestly from those of PEG. Since both PVP and PEG are flexible polymers with similar persistence lengths, their excluded volume contributions to μ_23_ are also expected to be similar. Since PEG is available at high purity over a much wider range of chain lengths than PVP, it is the better choice for these studies.

##### c) Applications of two-way alpha values to protein self-assembly interactions

Potential new directions of research using two-way alpha values include predicting or interpreting *χ* parameters of Flory-Huggins theory (59-61) in applications to aqueous polymer solutions. In this theory, *χ* quantifies the strength of segment-water interactions relative to segment-segment and water-water interactions. Extensions of this theory with “stickers and spacers” provide more realistic analyses of interactions of segments of flexible chain models of biopolymers (61, 62). It seem likely that two-way alpha values for amide atoms can be used to predict or interpret *χ* and noncoulombic “sticker” and “spacer” interaction parameters in analyses of the different behaviors of low-complexity polypeptides and unfolded proteins in chain expansion-collapse and aggregation (61, 63-65). Use of two-way alpha values would allow the sticker and spacer treatment to be extended to include a third type of region with net-unfavorable interactions (positive alpha values). Expansion of the set of two-way alpha values to include interactions of protein and nucleic acid unified atoms will allow their use in coarse-grained simulations and other analyses of interactions in liquid droplets formed by RNA and RNA binding proteins (61, 62, 65-67).

## Conclusion

Average strengths of interaction of amide O, N and C unified atoms, quantified per unit of accessible area of each atom by two-way alpha values, provide important bridges between protein structural (ASA) information, molecular dynamics simulations, and experimental studies of protein-solute interactions and solute effects of protein processes, as well as a window into a new chemistry of weak interactions of these O, N and C unified atoms in water.

## Acknowledgments

We thank Emily Zytkiewicz for comments on the manuscript, and gratefully acknowledge support of NIH GM R35-118100 for this research.

## Supplementary Information

### Materials and Methods

#### Chemicals

Formamide (>99.5%), N-methylformamide (>99%), and malonamide (>97%) were obtained from Sigma. Propionamide (>98%) was from Alfa Aesar and acetyl-L-ala-methylamide (aama, >99%) was from Bachem. All these amides were obtained in anhydrous form and used without further purification. All were dissolved in deionized water obtained from a Barnstead E-pure system (Thermo-Fischer Scientific).

#### Structures of Amide Compounds and ASA Calculations

Molecular structures of NEP (N-ethyl pyrrolidone) and of short oligomers of PVP (polyvinyl pyrrolidone) used for calculations of water-accessible surface areas (ASA) were predicted from NIH Cactus(1) website (https://cactus.nci.nih.gov/translate/) as described previously (2). Molecular structures of all other amide compounds investigated were analyzed previously (2). In all cases, a unified atom model was used in which hydrogens are treated as part of the C or N atom to which they are bonded. ASA information for NEP and PVP oligomers was calculated using the program Surface Racer (2) with the Richards set of van der Waals radii (3) and a 1.4 Å probe radius for water. As previously (2), ASA values were obtained for four coarse-grained atom types: amide sp^2^O, sp^2^N, sp^2^C and aliphatic sp^3^C. Alternative sources of structural information (PubChem ((4) and Biological Magnetic Resonance Bank (BMRB)(5)) and alternative ASA programs (VMD(6) and GetArea(7)) were compared with Cactus and Surface Racer previously (8, 9), and no significant differences were found.

#### Determination of ASA of End and interior Residues of PVP from Molecular Models of Short Oligomers

Water accessible surface areas (ASA) of the four types of unified atom (amide sp^2^O, sp^2^N, sp^2^C; aliphatic sp^3^C) of NEP and short PVP oligomers (number of residues N_3_ ≤ 5) were calculated using Surfracer program. Results are given in Table S8. To determine ASA contributions from the two end residues (*ASA*_2*E,i*_) and the N_3_-2 interior residues (*ASA*_*I,i*_) of a PVP oligomer, ASA values for each type of atom (i) were fitted to Eq.S1:

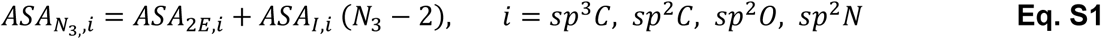

Values of *ASA*_2*E,i*_ and *ASA*_*I,i*_ obtained from these fits are also reported in Table S8.

#### Vapor Pressure Osmometry (VPO)

VPO is used to quantify thermodynamic interactions of small solutes which are soluble and nonvolatile in water by measuring osmolality differences Δ*Osm*(*m*_2_, *m*_3_) between three component (water, solute 2, and solute 3) and two component (water and solute 2, water and solute 3) solutions. Details of the osmolality analysis were described previously (9).

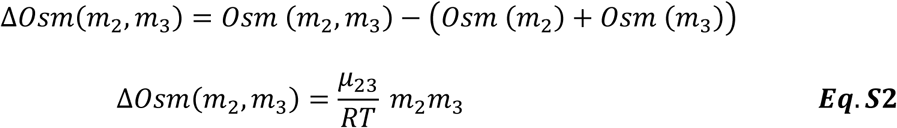

Preferential interactions (*μ*_23_ values) of a series of urea and alkyl ureas with one another and with other amide compounds were determined previously by VPO using Eq.S2 (9). Here *μ*_23_ values quantifying pairwise interactions in aqueous solution between five additional amide compounds (formamide, N-methylformamide, propionamide, malonamide and aama) are determined.

#### Analysis and Interpretation

##### One- and Two -Way Dissections of *μ*_23_ Values for Amide-Amide Interactions

In this section we provide specific expressions applying Eqs. 1-4 to the amides studied here.

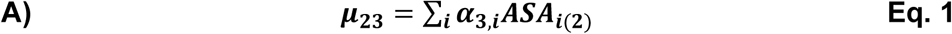

Each contribution in this sum is composed of an intrinsic interaction strength (one-way alpha value) for the interaction of solute *3* with a unit area (1 A^2^) of accessible surface of one type of unified atom of the biopolymer or other solute 2. For the interactions of amide compounds investigated here, these atom types are amide sp^2^O, sp^2^N and sp^2^C, and aliphatic sp^3^C. Taking as a specific example the interaction of acetyl-L-ala-methylamide (aama, component 3) with the various amide atoms of proprionamide (ppa, component 2), for which *μ*_23_= *μ*_*ppa,aama*_ = −43 ± 4 cal mol^-1^ molal^-1^ (Table S1):

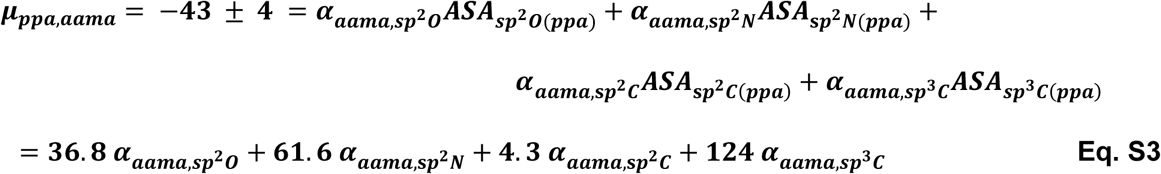

where the proprionamide ASA values in Eq. S3 (9) are in Å^2^ and the units of the one-way alpha values are cal mol^-1^ molal^-1^ Å^-2^.

Ten other equations like Eq. S3 are written to interpret experimental *μ*_2,*aama*_ values for the interactions of aama with formamide, N-methylformamide, malonamide, urea, methylurea, and the remainder of the set of amide compounds (component 2) investigated. Solving these eleven equations in four unknowns 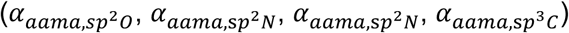 determines best-fit values for the above four one-way aama alpha values (Table S2). Comparison of predicted and observed *μ*_2,*aama*_ values for the full set of eleven aama-amide compound interactions (Fig. 2B, Table S3) tests the hypotheses of additivity and proportionality of contributions to ASA which are the basis of Eqs 1 and S3. Analogous sets of eleven equations are formulated and solved to obtain sets of four one-way alpha values quantifying the interactions of each other amide compound (formamide, N-methylformamide, malonamide, proprionamide) with a unit area of amide sp^2^O, sp^2^N, sp^2^C and aliphatic sp^3^C atoms (Table S2).

Since *μ*_23_= *μ*_32_ therefore

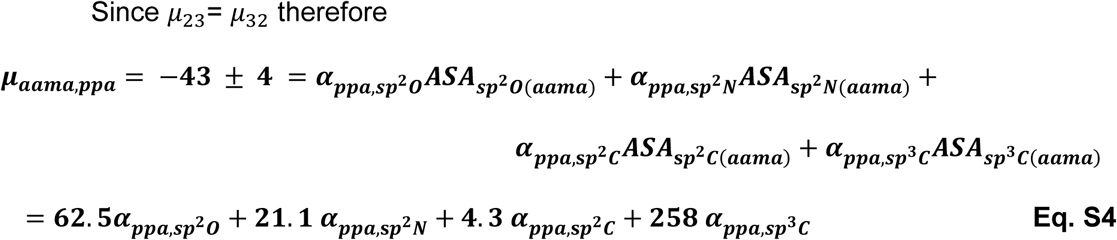

Ten other equations like Eq. S4 are written to interpret experimental *μ*_2,*ppa*_ values for the interactions of proprionamide with formamide, N-methylformamide, malonamide, urea, methylurea, and the remainder of the set of amide compounds (component 2) investigated.

Solving these eleven equations in four unknowns 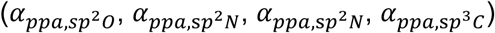 determines best-fit values for the above four one-way proprionamide alpha values (Table S2).

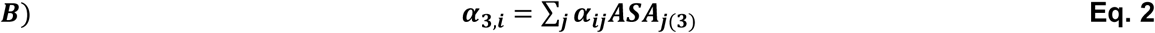

Continuing with the above example for the ppa-aama interaction,

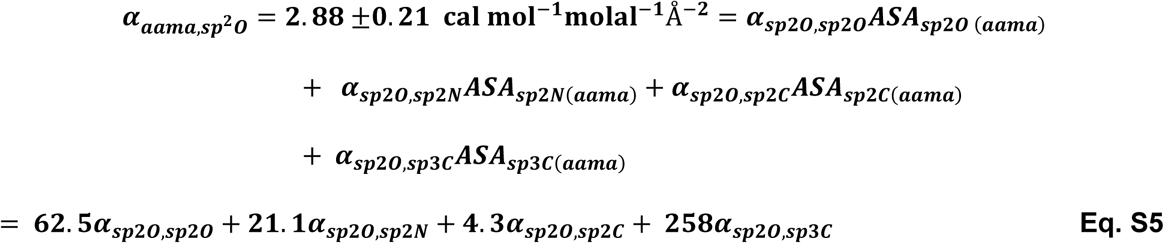

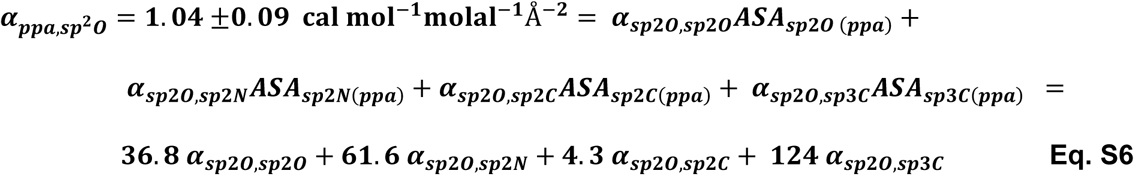

where the units of the two-way alpha values are cal mol^-1^ molal^-1^ Å^-4^. Nine other equations like Eqs. S5-6 are written to interpret experimental ***α***_**3,*sp*2*O***_ values for the interactions of formamide, N-methylformamide, malonamide, urea, methylurea, and the remainder of the set of amide compounds (component 2) with sp^2^O atoms. Solving these eleven equations in four unknowns (***α***_***Sp*2*O,Sp*2*O***_, ***α***_***Sp*2*O,Sp*2*N***_, ***α***_***Sp*2*O,Sp*2*C***_, ***α***_***Sp*2*O,Np*2*C***_) determines best-fit values for these four two-way alpha values (Table 1). Comparison of predicted and observed *μ*_23_ values for the full set of eleven aama-amide compound interactions (Fig. 2B, Table S3) tests the hypotheses of additivity and proportionality of contributions to ASA which are the basis of Eqs 2 and S3-6. An analogous set of eleven equations is formulated and solved to obtain a set of four two-way alpha values quantifying the interactions of a unit area of each other type of unified atom (amide sp^2^N, sp^2^C and aliphatic sp^3^C) with a unit area of amide sp^2^O, sp^2^N, sp^2^C and aliphatic sp^3^C atoms (Table 1).

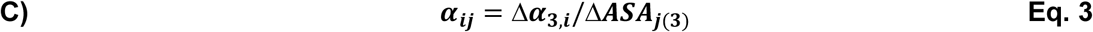

Here we illustrate the application of Eq. 3 to one of the four pairs of amides (methylurea and ethylurea) analyzed in the text. These amides differ primarily in amount of sp^3^C ASA. (See text for a discussion of all four amide pairs, based on the numerical analysis in Table S5.)

For methylurea (mu), the one-way α-value for the interaction with 1 Å^2^ of amide sp^2^O surface (0.78 cal mol^-1^ molal^-1^ Å^-2^; (9) is interpreted by Eq. 2 as the sum of ASA-weighted contributions from interactions of the methyl sp^3^C and amide sp^2^O, sp^2^N and sp^2^C atoms of methylurea (*mu*) with amide sp^2^O atoms of other compounds:

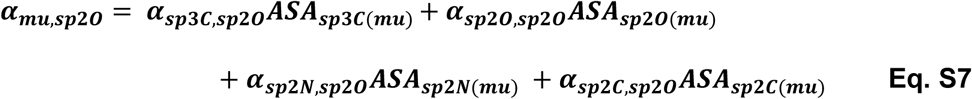

The corresponding equation for ethylurea (eu) is

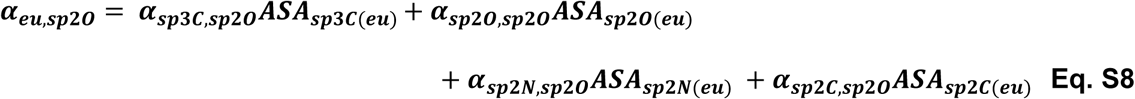

Subtracting Eq. S7 from S8 yields a specific example of Δ***α***_**3,*i***_:

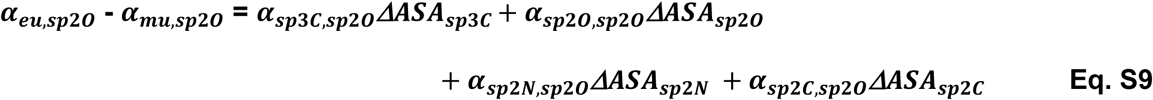

Because 87% of the ASA difference between ethylurea and methylurea is from sp^3^C, to a good approximation ***α***_***eu,Sp*2*O***_ **- *α***_***mu,sp*2*O***_ ≈ ***α***_***sp*3*C,sp*2*O***_***ΔASA*** _***sp*3*C***_ and

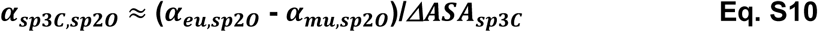

which is a specific example of Eq. 3 in the text. Table S5 summarizes the results of this and three other difference analyses to estimate two-way alpha values, and compares these estimates with those in Table 1, obtained from global fitting. For the case of ***α***_***Np*2*C,Sp*2*O***_ analyzed above, the estimate from Eq. S10 is within 30% of the Table 1 value, as shown in Table S5.

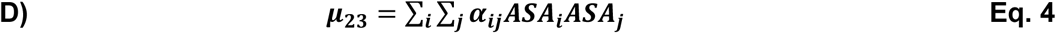

As in sections A-C above, indices i and j refer to the four types of unified atom present in the amide compounds investigated (amide sp^2^O, sp^2^N, sp^2^C; aliphatic sp^3^C). Hence, for each of the one hundred and five pairs of amide compounds investigated:

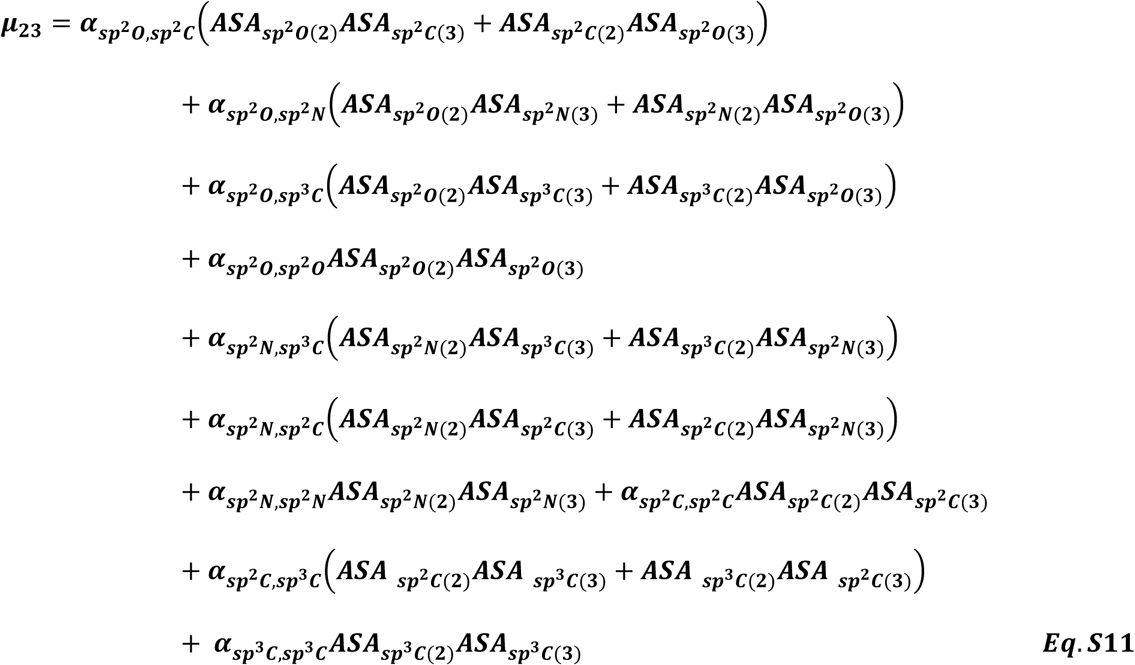

Here, as in sections A-C above, ASA_i(3)_ is the ASA of group i on solute 3 and ASA_j(2)_ is the ASA of group j on solute 2.

##### Predicting One-Way Alpha Values for Interactions of Amide Solutes with the Types of Unified Atoms of Amide Compounds

As described in the main text, two-way alpha values can be used to predict one way alpha values quantifying how any amide solute interacts with amide sp^2^O, sp^2^N, sp^2^C and aliphatic sp^3^C unified atoms on any amide or polyamide molecule or surface (e.g. the surface exposed in protein unfolding). As an example, one-way alpha values for interactions of all twelve amide solutes investigated with unit areas of the four amide unified atoms may be predicted from two-way alpha values (Table 1) and ASA information (9) using Eqs. 2 (see Eqs. S5-6 for examples), and compared with observed one-way alpha values determined from µ_23_ values using Eq. 1 (see Eqs. S3-4 for examples) and ASA information. Results of these two methods to obtain one-way alpha values are shown in Table S6. Agreement within the combined 1 SD uncertainties is observed for 83% of these solute-atom interactions, and all but the interaction of N-methylformamide with sp^2^O agree within 2 SD.

##### Comparison of Two-way Alpha Values for Atom-Atom Interactions of Amides From Different Treatments of sp^2^C

One-way alpha values for interactions of urea and alkylureas with amide and aromatic sp^2^C were found to be similar (9). Two-way alpha values listed in Table 1 were determined by analysis of 105 µ_23_ values for amide interactions (85 amide compound-amide compound, 20 amide compound-aromatic compound) using Eq. 4, to obtain a combined two-way alpha value for amide and aromatic sp^2^C. To justify this analysis, here we extend it by global fitting all µ_23_ values (105 total, including 20 for amide-aromatic hydrocarbon interactions) to Eq. S12 which includes one global weighting factor 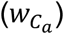 quantifying the relative strength of interactions of amide sp^2^C as compared to aromatic sp^2^C. Clearly this is an oversimplification, since in principle a different weighting factor might be needed for interactions of sp^2^C with each other type of atom, but it provides a test of whether such corrections are significant. The revised version of Eq. 4 for *μ*_23_ is

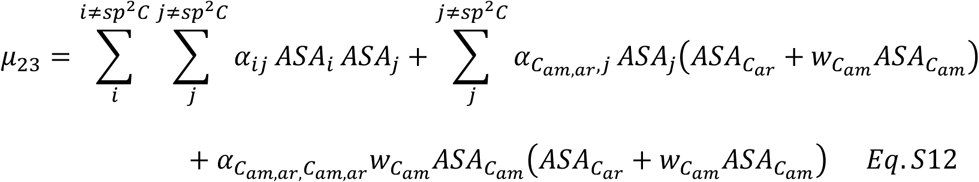

In Eq. S12, the subscript C_am,ar_ stands for combined amide and aromatic sp^2^C, C_am_ stands for amide sp^2^C and C_ar_ is aromatic sp^2^C.

Two-way alpha values summarized in Table 1 were obtained for the unweighted case 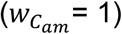. In Table S7 these values are compared with those obtained from a global analysis using Eq. S12 and floating 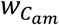. Two-way alpha values obtained from this analysis are the same as in Table 1 within the uncertainty, although the percent difference in the interaction of sp^2^O with sp^2^O is about 80%. In this fit, the weighting coefficient 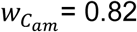, indicating that on average interactions of amide sp^2^C with the different atom types are about 82% as strong as for aromatic sp^2^C.

Table S7 also compares two-way alpha values obtained from analyses of subsets of *μ*_23_ values with those in Table 1 and from the fit with 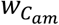 weighting. Fitting only the 64 amide-amide *μ*_23_ values obtained for amides with minimal amounts of amide sp^2^C to the variant of Eq. 4 with only six terms for interactions involving only sp^2^O, sp^2^N and sp^3^C yields six two-way alpha values which agree within the combined uncertainty with those of Table 1. Fitting only the 20 amide-aromatic *μ*_23_ values to another variant of Eq. 4 yields two-way alpha values for interactions of sp^2^C with sp^2^O, sp^2^N, sp^2^C and sp^3^C. Two-way alpha values for sp^2^C-sp^2^C and sp^2^C-sp^3^C agree with those in Table 1, while those for sp^2^C-sp^2^O, and sp^2^C-sp^2^N are both 20-30% larger in magnitude than their counterparts in Table 1, consistent with the finding of a weighting coefficient 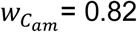 for interactions involving amide sp^2^C.

##### Additional Tests of Effect of Size of Dataset on Two-way Alpha Values

Table S7, discussed above, compared the separate determinations of four two-way alpha values from *μ*_23_ values for 20 amide-aromatic interactions and of six two-way alpha values for 85 amide-amide interactions with the ten two-way alpha values obtained from global analysis of the set of 105 *μ*_23_ values for amide-aromatic and amide-amide interactions, treating amide sp^2^C as the same as or differently from aromatic sp^2^C. Two-way alpha values obtained from these various approaches agree in most cases within the combined uncertainty. In Table S11 the effect of other reductions in the size of the *μ*_23_ data set are examined. This table shows there is little effect on two-way alpha values of removing all of the 14 to 26 *μ*_23_ values that quantify interactions of the more polar (urea, malonamide) and/or nonpolar (,3-diethylurea, aama) amides. The insensitivity of the two-way alpha values to these reductions in the set of *μ*_23_ values analyzed shows that even these subsets are large enough and diverse enough to determine all ten two-way alpha values.

##### Predicting the Chemical Interactions of PVP and its Model Monomer NEP with Amide and Hydrocarbon Atoms of Proteins

This section generalizes Eq. 6 for chemical (preferential interaction) contributions to *μ*_23_ values for interactions of PVP oligomers or polymers of any number of residues (N_3_) with the different hybridization states of O, N and C atoms of other solutes or proteins. For interactions of larger PVP oligomers and polymers with large solutes, an excluded volume term also contributes to *μ*_23_ and the chemical term in Eq. 6 may be reduced by a shielding term χ (10). Since any PVP has two end residues and N_3_-2 interior residues, the interaction of the average PVP residue with a solute 2 is therefore

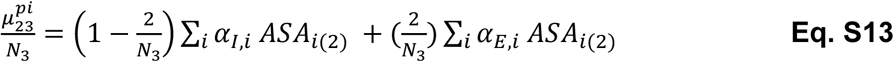

For N_3_ > 20, Eq. S13 reduces to Eq. 6 of the main text. In Eq. S13, *α*_*E,i*_ and *α*_*I,i*_ are one-way alpha values that quantify the intrinsic strength of interactions of PVP end (E) and interior (I) residues with the i-th type of atom on another solute or protein. For PEG, where the end residues (as defined) are half the size of interior residues, we combined them (*α*_2*E,i*_) but for PVP it is more appropriate to treat each end residue separately. In Eq. 6 of the text for *μ*_23_ for high molecular weight PVP (N_3_ >> 1)), no distinction is made between end and interior residues.

By analogy with Eq. 2, each one-way *α*_*E,i*_ and *α*_*I,i*_ in Eq. S13 is itself a sum of contributions of interaction of the i-th type of protein atom with the j-th type of PVP atom (see Eq.3).

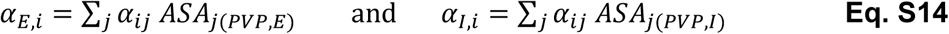

In Eq. S14, each two-way alpha value (*α*_*ii*_) quantifies the interaction of 1 A^2^ atom type i of solute 2 with 1 A^2^ of PVP atom type j, *ASA*_*i*(*PVP,E*)_ and *ASA*_*i*(*PVP,I*)_ are areas (in A^2^) of atom type j on the end and interior residues of PVP. One-way alpha values for NEP and for PVP end and interior residues, calculated from two-way alpha values as in Eqs. 2 and S14, are listed in Table S9 and compared with the corresponding quantities for PEG.

As previously (10), we interpret the interaction of PVP (component 3) with a protein (component 2) as the sum of preferential interaction (abbreviated pi) and excluded volume (ev) contributions,

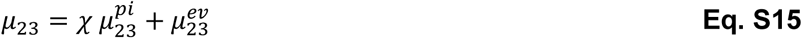

In Eq. S15, the quantity *χ* is the fractional accessibility of the average residue of PVP. For a PVP oligomer one expects *χ* ≈1, but for polymeric PVP one expects *χ* ≪ 1. (10)

**Figure S1.**
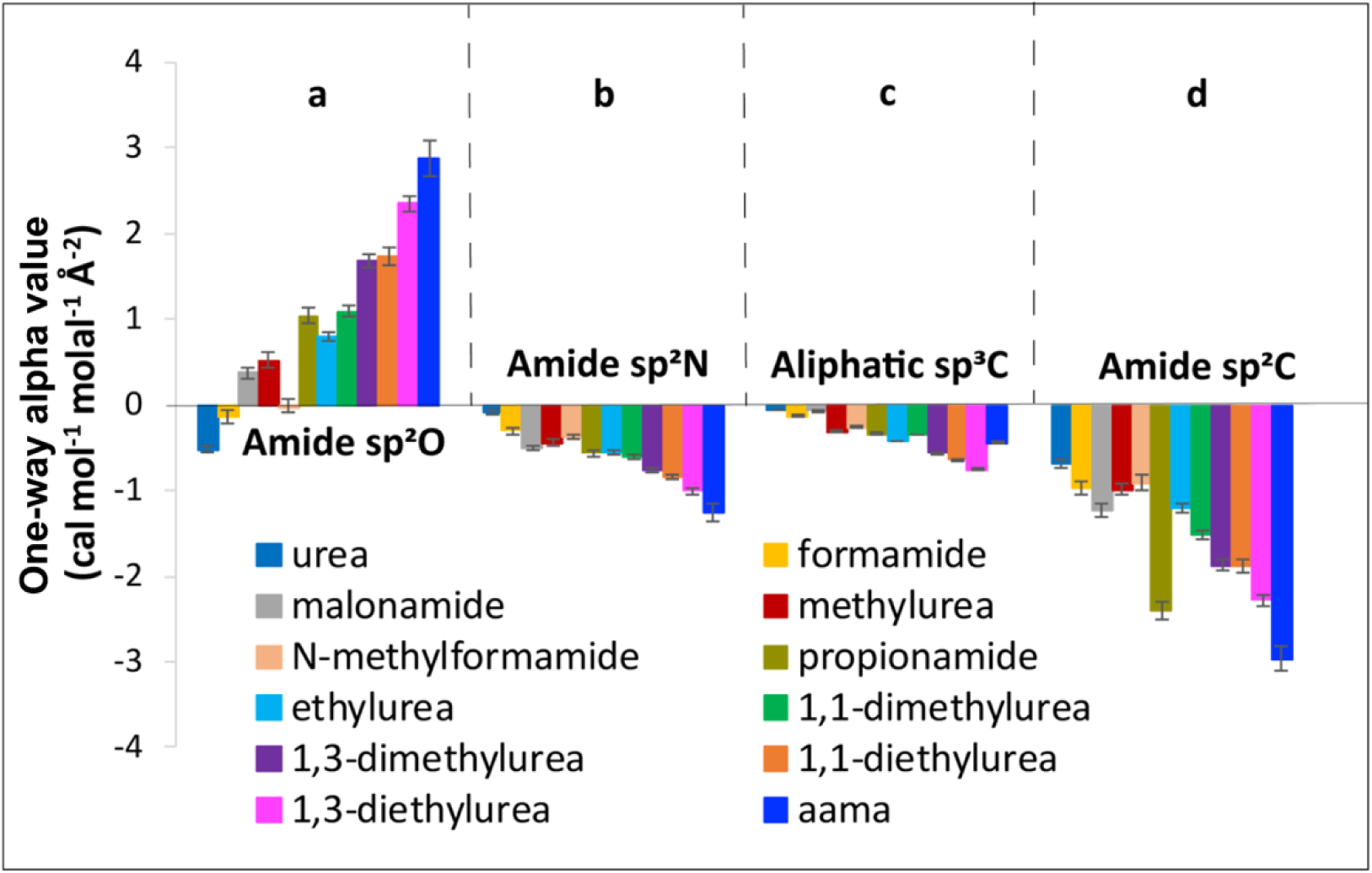
Comparison of One Way Alpha Values for Formamide, N-methyl Formamide, Malonamide, Proprionamide and N-acetylalanine N-methylamide (aama) with Other Amides. Amide compounds are listed arbitrarily in order of increasing aliphatic sp^3^C + amide sp^2^C ASA. Bar graphs compare interaction potentials (One-way alpha values; Table S1)) quantifying interactions of each amide compound with a unit area of a) amide sp^2^O, b) amide sp^2^N, c) amide-context sp^3^C, and d) amide/aromatic sp^2^C at 23 °C. Favorable interactions have negative α-values while unfavorable interactions have positive α-values. (aama: N-acetylalanine N-methylamide) α-Values for urea and alkyl ureas were reported previously (9).

**Figure S2.**
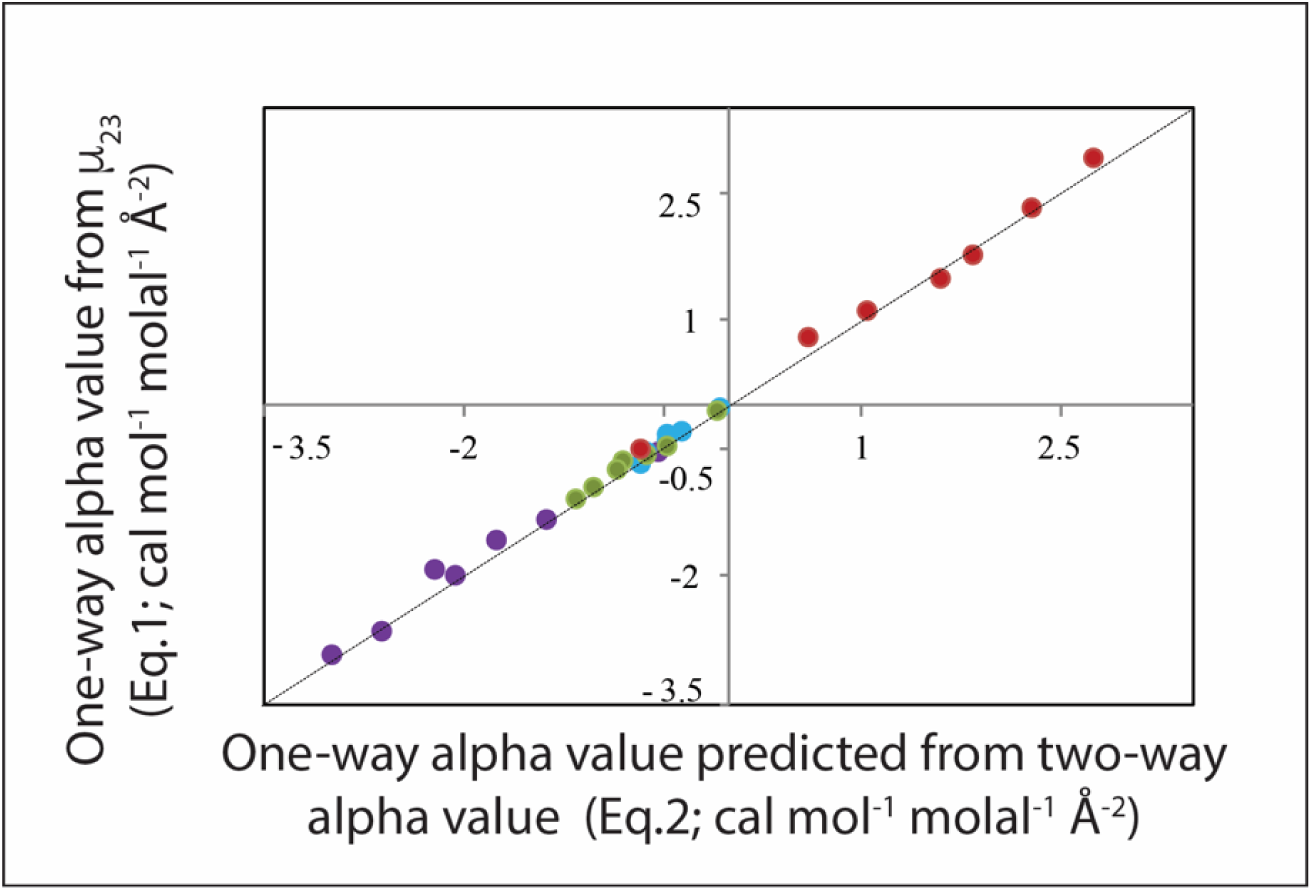
Comparison of Predicted and Observed one-way alpha values (cal mol^-1^ m^-1^ Å^-2^) for Interactions of Urea and Alkylureas (urea, methylurea, ethylurea, 1,1-dimethylrea, 1,3-dimethylurea, 1,1-diethylurea and/or 1,3-diethylurea) with amide and aromatic functional groups (amide sp^2^O, amide sp^2^N, amide sp^3^C and combined aromatic and amide sp^2^C) at 23 °C; Observed one-way α_i_-values of these ureas were determined in reference (9) and predictions of one-way α-values use two-way alpha values of amide-amide interactions (Table 1) and the values are also tabulated in Table S6 above. The line represents equality of predicted and observed values.

**Figure S3.**
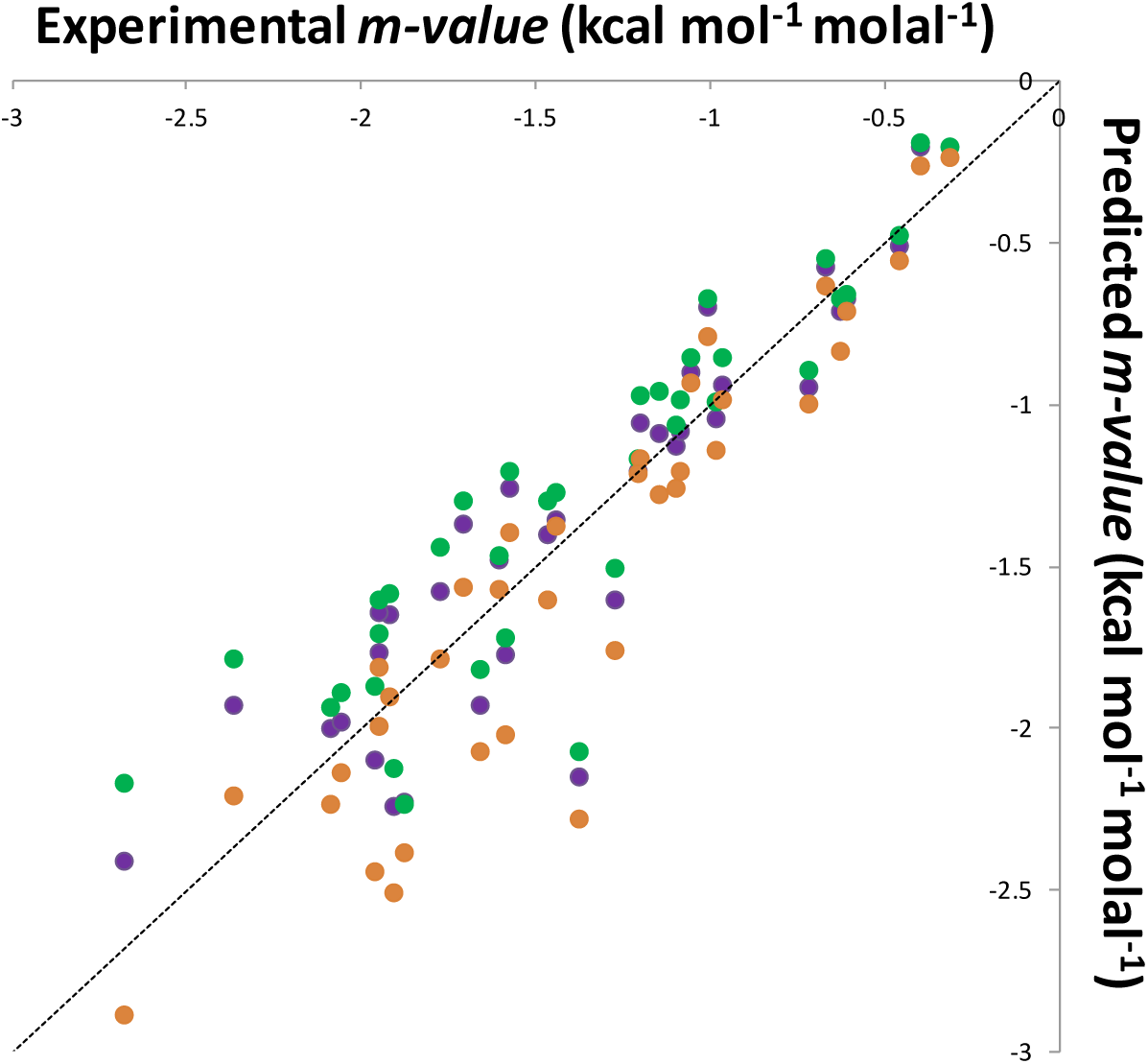
Comparison of Predicted and Observed Urea *m*-values of Unfolding Globular Proteins. ΔASA Values for unfolding of these proteins are from reference (11). One-way alpha values of urea with 7 protein functional groups are from reference (12). One-way alpha values of urea with four unified atoms of amides were determined in reference (9); Two-way alpha values of atom-atom interactions are determined in this work (α_ij_-values in Table 1). Purple: Previously-reported predictions of urea *m*-values using seven urea α-values (including hydroxyl O, carboxylate O and cationic N in addition to above amide and hydrocarbon unified atoms); Green: Predicted *m*-values were obtained from only four urea α-values (aromatic sp^2^C, aliphatic sp^3^C, amide sp^2^O and amide sp^2^N; reference (9). Yellow: Predicted *m*-values use two-way alpha values (Table 1) using Eq.3. Amide sp^2^C represents less than 1% of the ΔASA of unfolding and was not accounted for in these comparisons.

### Supplemental Tables

**Table S1.** Interactions of Amides with Relatively Large sp^2^C and/or sp^2^O Surface Area.

| Amide Compounds | | Observed $\mu_{23}$ value<sup>a</sup> |
| --- | --- | --- |
| propionamide | N-acetylalanine-N-methylamide | $-43.2 \pm 3.9$ |
| propionamide | formamide | $-76.8 \pm 2.6$ |
| propionamide | N-methylformamide | $-102.3 \pm 1.9$ |
| malonamide | N-acetylalanine-N-methylamide | $-8.6 \pm 2.2$ |
| malonamide | formamide | $-54.1 \pm 2.4$ |
| malonamide | N-methylformamide | $-64.5 \pm 3.0$ |
| N-acetylalanine-N-methylamide | formamide | $-53.0 \pm 3.1$ |
| N-acetylalanine-N-methylamide | N-methylformamide | $-80.3 \pm 3.2$ |
| formamide | N-methylformamide | $-64.0 \pm 2.2$ |

<sup>a</sup> Values of $\mu_{23} = \mu_{32}$ determined by VPO at 23°C. Units of $\mu_{23}$ are $\text{cal mol}^{-1} \text{molal}^{-1}$ .

**Table S2.** One-way Alpha Values for Interactions of Formamide, Malonamide, N-methylformamide, Propionamide and aama^a^ With Amide sp^2^O, sp^2^N, sp^2^C and Aliphatic sp^3^C Atoms.

|  | Amide sp<sup>2</sup>O | Amide sp<sup>2</sup>N | Aliphatic sp<sup>3</sup>C | Amide sp<sup>2</sup>C |
| --- | --- | --- | --- | --- |
| formamide | -0.13 ± 0.08 | -0.33 ± 0.03 | -0.14 ± 0.01 | -0.92 ± 0.08 |
| malonamide | 0.37 ± 0.07 | -0.5 ± 0.02 | -0.07 ± 0.01 | -1.23 ± 0.07 |
| N-methylformamide | -0.02 ± 0.09 | -0.41 ± 0.03 | -0.27 ± 0.01 | -0.81 ± 0.09 |
| propionamide | 1.04 ± 0.09 | -0.56 ± 0.04 | -0.33 ± 0.01 | -2.39 ± 0.1 |
| aama<sup>a</sup> | 2.88 ± 0.21 | -1.26 ± 0.09 | -0.45 ± 0.01 | -2.97 ± 0.14 |

<sup>a</sup> aama: N-acetylalanine-N-methylamide.
<sup>b</sup> One-way alpha values are obtained by fitting 11 experimental $\mu_{23}$ values for each compound listed (Tables S1, S3) to Eq.1. Uncertainties in alpha values are calculated as previously described.(13)

**Table S3.**
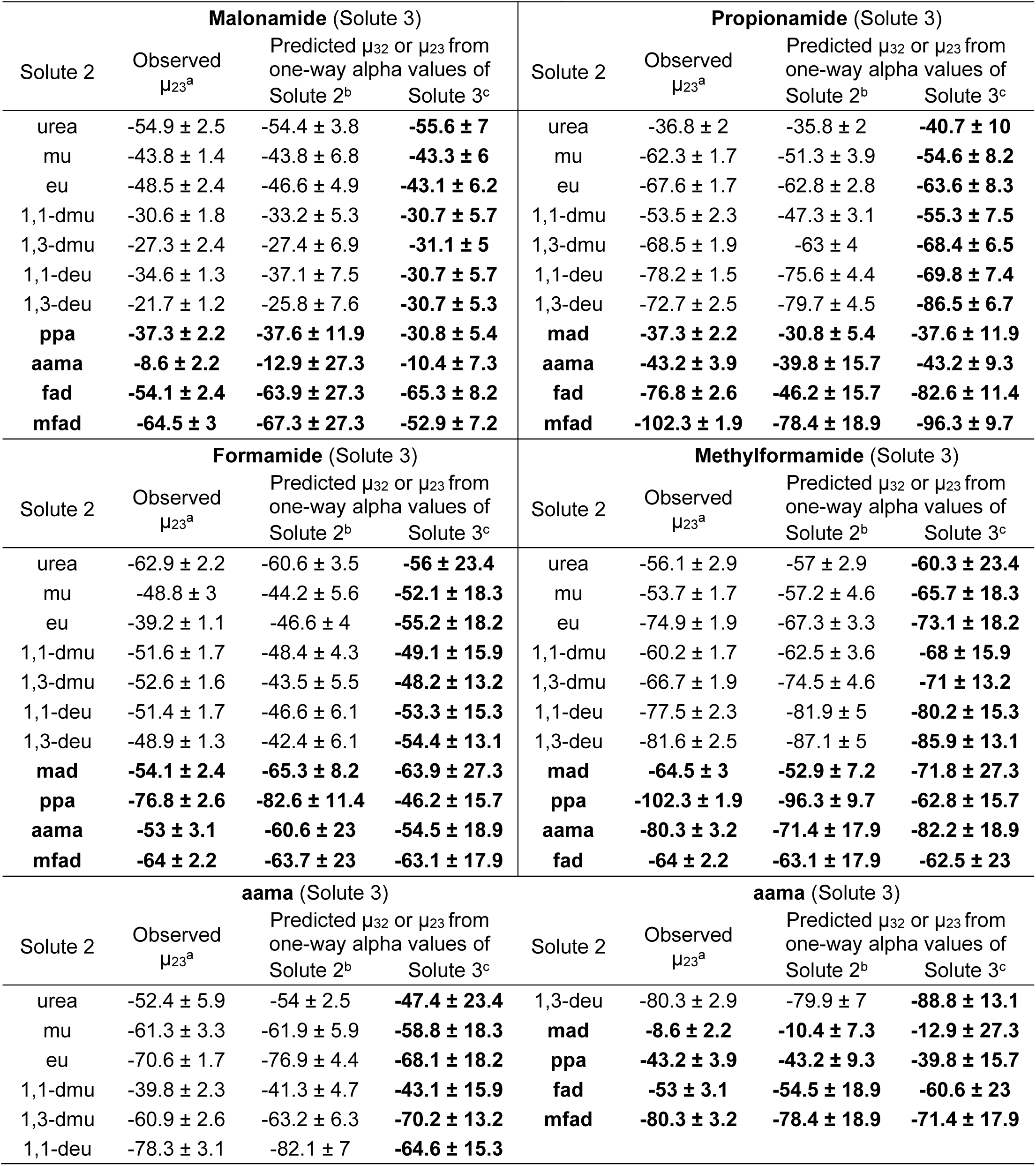
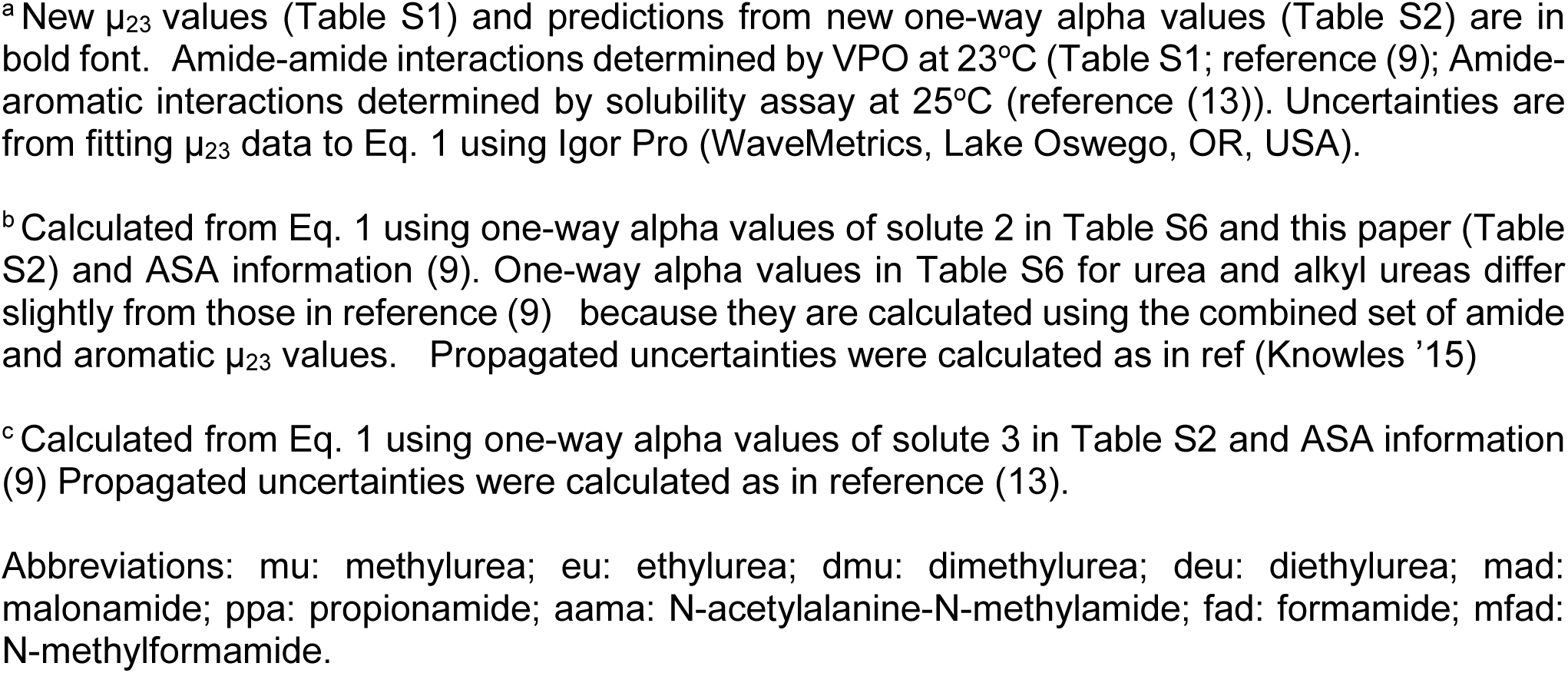
Comparison of Observed µ_23_ Values (cal mol^-1^ molal^-1^; µ_23_ = µ_32_) for Amide Interactions at 23°C with Predictions of µ_23_ and µ_32_ from One-way Alpha Values^a^.

**Table S4.**
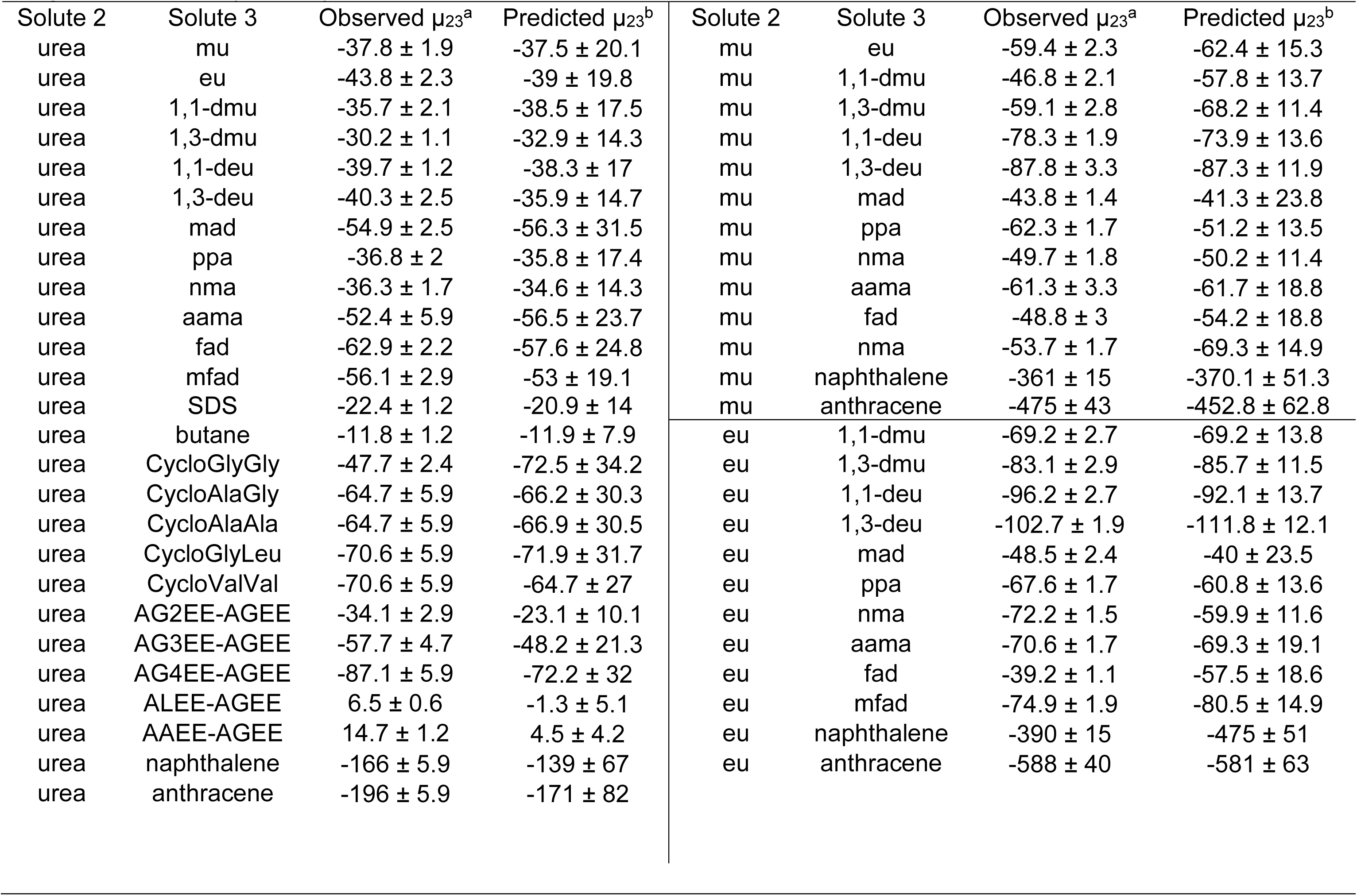
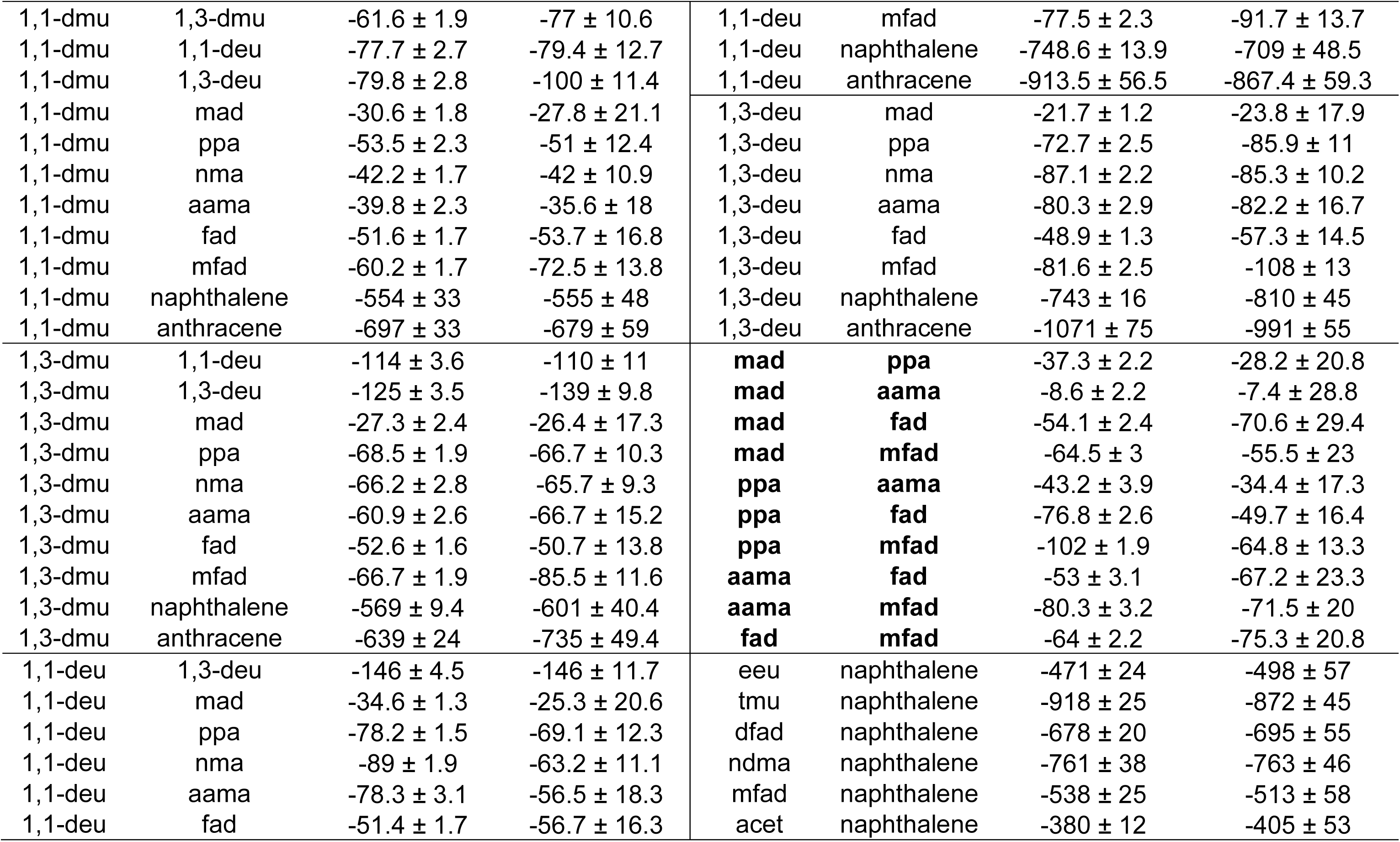
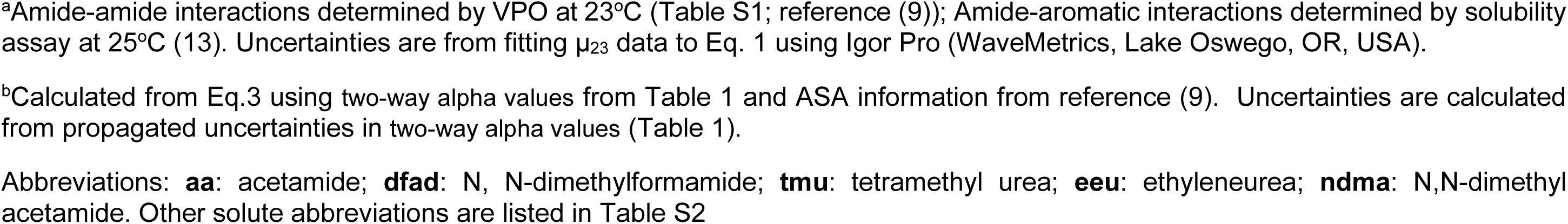
Comparison of Observed µ_23_ Values (cal mol^-1^ molal^-1^) for Amide Interactions with Predictions from Two-Way Alpha Values (Table 1)

**Table S5.** Comparison of Two-way Alpha Values Obtained by Fitting with Estimates from One-way Alpha Values for Amide Pairs Differing Primarily in ASA of One Atom Type.

| Amide Pair<sup>a</sup> | %ΔASA<sup>b</sup> | Two-way Alpha Value (cal mol<sup>-1</sup> molal<sup>-1</sup> Å<sup>-4</sup>) |  |  |  |  |  |  |  |
| --- | --- | --- | --- | --- | --- | --- | --- | --- | --- |
|  |  | sp<sup>2</sup>O |  | sp<sup>2</sup>N |  | sp<sup>2</sup>C |  | sp<sup>3</sup>C |  |
|  |  | Estimate<br>(Eq. 3) | Fitted<br>(Table 1) | Estimate<br>(Eq. 3) | Fitted<br>(Table 1) | Estimate<br>(Eq. 3) | Fitted<br>(Table 1) | Estimate<br>(Eq. 3) | Fitted<br>(Table 1) |
| eu–mu | (86% sp<sup>3</sup>C)<sup>c</sup> | 0.008 | 0.011 | -0.003 | -0.0038 | -0.006 | -0.010 | -0.003 | -0.0039 |
| 1,1-deu – 1,1-dmu | (90% sp<sup>3</sup>C)<sup>d</sup> | 0.010 | 0.011 | -0.004 | -0.0038 | -0.006 | -0.010 | -0.004 | -0.0039 |
| 1,3-deu – ppa | (87% sp<sup>2</sup>N)<sup>e</sup> | -0.012 | -0.011 | 0.001 | 0.0034 | (0.058) | 0.0018 | -0.004 | -0.0038 |
| aama – 1,3-deu | (63% sp<sup>2</sup>O)<sup>f</sup> | 0.016 | 0.018 | -0.009 | -0.011 | -0.020 | -0.014 | 0.009 | 0.011 |

<sup>a</sup> Abbreviations for amide compounds as in Table S3-4.
<sup>b</sup> ΔASA calculated as the sum of magnitudes of ASA differences for the different atom types.
<sup>c</sup> ΔASA = 42 Å<sup>2</sup> (36 Å<sup>2</sup> sp<sup>3</sup>C, -6 Å<sup>2</sup> sp<sup>2</sup>N)
<sup>d</sup> ΔASA = 71 Å<sup>2</sup> (64 Å<sup>2</sup> sp<sup>3</sup>C, -4 Å<sup>2</sup> sp<sup>2</sup>N, -3 Å<sup>2</sup> sp<sup>2</sup>O)
<sup>e</sup> ΔASA = 23 Å<sup>2</sup> (20 Å<sup>2</sup> sp<sup>2</sup>N, 2 Å<sup>2</sup> sp<sup>3</sup>C, 1 Å<sup>2</sup> sp<sup>2</sup>O)
<sup>f</sup> ΔASA = 54 Å<sup>2</sup> (34 Å<sup>2</sup> sp<sup>2</sup>O, -13 Å<sup>2</sup> sp<sup>2</sup>N; 7 Å<sup>2</sup> sp<sup>3</sup>C)

**Table S6.** Comparison of One-way Alpha Values (cal mol^-1^ molal^-1^ Å^-2^) Calculated from µ_23_ Values with Predictions of One-way Alpha Values from Two-way Alpha Values.

| Solute <sup>a</sup> | One-way Alpha Value (cal mol<sup>-1</sup> m<sup>-1</sup> Å<sup>-2</sup>) |  |  |  |  |  |  |  |
| --- | --- | --- | --- | --- | --- | --- | --- | --- |
|  | Amide sp<sup>2</sup>O <sup>b</sup> |  | Amide sp<sup>2</sup>N <sup>b</sup> |  | Amide sp<sup>3</sup>C <sup>b</sup> |  | Aromatic/Amide sp<sup>2</sup>C <sup>b</sup> |  |
| | Calculated from $\mu_{23}$ <sup>c</sup> | Predicted <sup>d</sup> | Calculated from $\mu_{23}$ <sup>c</sup> | Predicted <sup>d</sup> | Calculated from $\mu_{23}$ <sup>c</sup> | Predicted <sup>d</sup> | Calculated from $\mu_{23}$ <sup>c</sup> | Predicted <sup>d</sup> |
| urea | -0.53 ± 0.03 | -0.64 ± 0.33 | -0.1 ± 0.02 | -0.06 ± 0.17 | -0.06 ± 0 | -0.05 ± 0.05 | -0.59 ± 0.01 | -0.51 ± 0.37 |
| mu | 0.78 ± 0.08 | 0.61 ± 0.25 | -0.51 ± 0.03 | -0.44 ± 0.14 | -0.34 ± 0.01 | -0.33 ± 0.04 | -1.36 ± 0.08 | -1.36 ± 0.32 |
| eu | 1.08 ± 0.07 | 1.06 ± 0.25 | -0.62 ± 0.02 | -0.59 ± 0.15 | -0.46 ± 0.01 | -0.45 ± 0.05 | -1.6 ± 0.07 | -1.74 ± 0.33 |
| 1,1-dmu | 1.45 ± 0.07 | 1.63 ± 0.23 | -0.69 ± 0.03 | -0.78 ± 0.13 | -0.39 ± 0.01 | -0.43 ± 0.04 | -2.03 ± 0.07 | -2.03 ± 0.3 |
| 1,3-dmu | 1.74 ± 0.06 | 1.86 ± 0.19 | -0.78 ± 0.03 | -0.82 ± 0.11 | -0.57 ± 0.01 | -0.61 ± 0.04 | -1.95 ± 0.04 | -2.2 ± 0.26 |
| 1,1-deu | 2.31 ± 0.09 | 2.31 ± 0.23 | -0.98 ± 0.03 | -1 ± 0.13 | -0.7 ± 0.01 | -0.66 ± 0.04 | -2.69 ± 0.1 | -2.6 ± 0.31 |
| 1,3-deu | 2.89 ± 0.12 | 2.77 ± 0.21 | -1.14 ± 0.04 | -1.13 ± 0.12 | -0.83 ± 0.02 | -0.85 ± 0.04 | -2.97 ± 0.13 | -2.97 ± 0.28 |
| Solute | Amide sp<sup>2</sup>O <sup>e</sup> |  | Amide sp<sup>2</sup>N <sup>e</sup> |  | Amide sp<sup>3</sup>C <sup>e</sup> |  | Amide sp<sup>2</sup>C <sup>e</sup> |  |
| | Calculated from $\mu_{23}$ <sup>c</sup> | Predicted <sup>d</sup> | Calculated from $\mu_{23}$ <sup>c</sup> | Predicted <sup>d</sup> | Calculated from $\mu_{23}$ <sup>c</sup> | Predicted <sup>d</sup> | Calculated from $\mu_{23}$ <sup>c</sup> | Predicted <sup>d</sup> |
| fad | -0.13 ± 0.08 | -0.4 ± 0.32 | -0.33 ± 0.03 | -0.24 ± 0.18 | -0.14 ± 0.01 | -0.13 ± 0.06 | -0.92 ± 0.08 | -1.03 ± 0.42 |
| mfad | -0.02 ± 0.09 | 0.86 ± 0.27 | -0.41 ± 0.03 | -0.62 ± 0.15 | -0.27 ± 0.01 | -0.4 ± 0.05 | -0.81 ± 0.09 | -1.88 ± 0.36 |
| mad | 0.37 ± 0.07 | 0.26 ± 0.4 | -0.5 ± 0.02 | -0.46 ± 0.21 | -0.07 ± 0.01 | -0.03 ± 0.06 | -1.23 ± 0.07 | -1.28 ± 0.45 |
| ppa | 1.04 ± 0.09 | 1.29 ± 0.22 | -0.56 ± 0.04 | -0.65 ± 0.13 | -0.33 ± 0.01 | -0.36 ± 0.04 | -2.39 ± 0.1 | -1.73 ± 0.29 |
| aama | 2.88 ± 0.21 | 3.63 ± 0.35 | -1.26 ± 0.09 | -1.57 ± 0.17 | -0.45 ± 0.01 | -0.45 ± 0.06 | -2.97 ± 0.14 | -3.53 ± 0.4 |

<sup>a</sup> See Table S3-4 for abbreviations.
<sup>b</sup> Calculated from $\mu_{23}$ values for amide-amide and amide-aromatic interactions assuming no distinction between aromatic sp<sup>2</sup>C and amide sp<sup>2</sup> C (see Table S7 reference (9)).
<sup>c</sup> Calculated from Eq. 1.
<sup>d</sup> Predicted from two-way alpha values and ASA information using Eq. 2.
<sup>e</sup> Calculated from $\mu_{23}$ values for amide-amide interactions only.

**Table S7.**
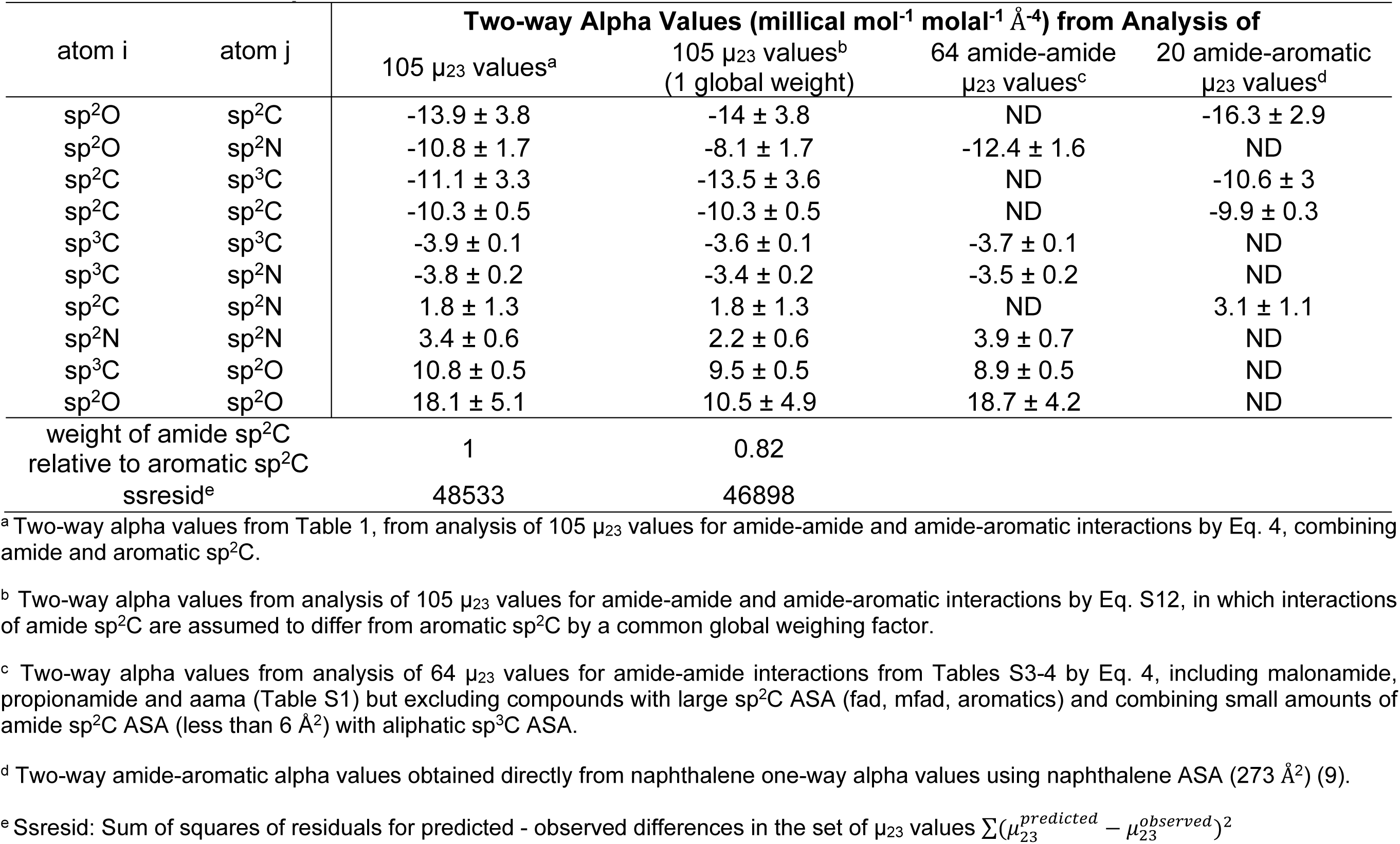
Comparison of Two-way Alpha Values for Atom-atom Interactions Using Different Treatments of Amide and Aromatic sp^2^C.

| atom i | atom j | Two-way Alpha Values (millical mol<sup>-1</sup> molal<sup>-1</sup> Å<sup>-4</sup>) from Analysis of |  |  |  |
| --- | --- | --- | --- | --- | --- |
|  |  | 105 μ<sub>23</sub> values<sup>a</sup> | 105 μ<sub>23</sub> values<sup>b</sup><br>(1 global weight) | 64 amide-amide<br>μ<sub>23</sub> values<sup>c</sup> | 20 amide-aromatic<br>μ<sub>23</sub> values<sup>d</sup> |
| sp<sup>2</sup>O | sp<sup>2</sup>C | -13.9 ± 3.8 | -14 ± 3.8 | ND | -16.3 ± 2.9 |
| sp<sup>2</sup>O | sp<sup>2</sup>N | -10.8 ± 1.7 | -8.1 ± 1.7 | -12.4 ± 1.6 | ND |
| sp<sup>2</sup>C | sp<sup>3</sup>C | -11.1 ± 3.3 | -13.5 ± 3.6 | ND | -10.6 ± 3 |
| sp<sup>2</sup>C | sp<sup>2</sup>C | -10.3 ± 0.5 | -10.3 ± 0.5 | ND | -9.9 ± 0.3 |
| sp<sup>3</sup>C | sp<sup>3</sup>C | -3.9 ± 0.1 | -3.6 ± 0.1 | -3.7 ± 0.1 | ND |
| sp<sup>3</sup>C | sp<sup>2</sup>N | -3.8 ± 0.2 | -3.4 ± 0.2 | -3.5 ± 0.2 | ND |
| sp<sup>2</sup>C | sp<sup>2</sup>N | 1.8 ± 1.3 | 1.8 ± 1.3 | ND | 3.1 ± 1.1 |
| sp<sup>2</sup>N | sp<sup>2</sup>N | 3.4 ± 0.6 | 2.2 ± 0.6 | 3.9 ± 0.7 | ND |
| sp<sup>3</sup>C | sp<sup>2</sup>O | 10.8 ± 0.5 | 9.5 ± 0.5 | 8.9 ± 0.5 | ND |
| sp<sup>2</sup>O | sp<sup>2</sup>O | 18.1 ± 5.1 | 10.5 ± 4.9 | 18.7 ± 4.2 | ND |
| weight of amide sp<sup>2</sup>C<br>relative to aromatic sp<sup>2</sup>C |  | 1 | 0.82 |  |  |
| ssresid<sup>e</sup> |  | 48533 | 46898 |  |  |

<sup>a</sup> Two-way alpha values from Table 1, from analysis of 105 μ<sub>23</sub> values for amide-amide and amide-aromatic interactions by Eq. 4, combining amide and aromatic sp<sup>2</sup>C.
<sup>b</sup> Two-way alpha values from analysis of 105 μ<sub>23</sub> values for amide-amide and amide-aromatic interactions by Eq. S12, in which interactions of amide sp<sup>2</sup>C are assumed to differ from aromatic sp<sup>2</sup>C by a common global weighing factor.
<sup>c</sup> Two-way alpha values from analysis of 64 μ<sub>23</sub> values for amide-amide interactions from Tables S3-4 by Eq. 4, including malonamide, propionamide and aama (Table S1) but excluding compounds with large sp<sup>2</sup>C ASA (fad, mfad, aromatics) and combining small amounts of amide sp<sup>2</sup>C ASA (less than 6 Å<sup>2</sup>) with aliphatic sp<sup>3</sup>C ASA.
<sup>d</sup> Two-way amide-aromatic alpha values obtained directly from naphthalene one-way alpha values using naphthalene ASA (273 Å<sup>2</sup>) (9).
<sup>e</sup> Ssresid: Sum of squares of residuals for predicted - observed differences in the set of μ<sub>23</sub> values $\sum(\mu_{23}^{predicted} - \mu_{23}^{observed})^2$

**Table S8.** PVP Oligomer ASA Values in Å^2^ from Cactus Models; Comparison with PubChem.

| Number of oligomer residues N<sub>3</sub> | Source | Amide sp<sup>2</sup>O | Amide sp<sup>2</sup>N | Aliphatic sp<sup>3</sup>C | Amide sp<sup>2</sup>C | Total |
| --- | --- | --- | --- | --- | --- | --- |
| 1 (NEP)<sup>a</sup> | Cactus | 36.9 | 0.35 | 252 | 2.6 | 292 |
| 2<sup>a</sup> |  | 54.5 | 0.35 | 381 | 3.0 | 439 |
| 3<sup>a</sup> |  | 81.5 | 0.35 | 479 | 3.4 | 564 |
| 4<sup>a</sup> |  | 88.9 | 0.35 | 578 | 3.8 | 671 |
| 5<sup>a</sup> |  | 97.5 | 0.35 | 674 | 3.4 | 775 |
| Interior residue<sup>b</sup> | Cactus (linear fitting) | 15.6 | 0 | 104 | 0.2 | 120 |
| End residues<sup>b</sup> |  | 56.3 | 0.35 | 369 | 3.0 | 428 |
| 1 (NEP) | PubChem | 39.0 | 0.52 | 250 | 2.8 | 292 |
| 2<sup>c</sup> |  | 54.6 | 0.35 | 381 | 3.0 | 439 |

<sup>a</sup> Molecular models created as described in SI text. Chemical formula of interior PVP residue is C<sub>6</sub>H<sub>9</sub>NO, end residue C<sub>6</sub>H<sub>10</sub>NO
<sup>b</sup> Determined with Eq. S1 as described in SI

**Table S9.** PVP One-Way End and Interior Residue Alpha Values; Comparison with NEP and PEG.

| Protein Atom Type | NEP (Monomer)<br>($\alpha_{NEP,i}$)<sup>a</sup> | PVP Interior Residue ($\alpha_{I,i}$)<sup>a</sup> | PEG Interior Residue ($\alpha_{I,i}$)<sup>a</sup> | PVP End Residues ($\alpha_{E,i}$)<sup>a</sup> | PEG End Residues ($\alpha_{2E,i}$)<sup>a</sup> |
| --- | --- | --- | --- | --- | --- |
| Aliphatic sp<sup>3</sup>C | 0.61 ± 0.03 | -0.24 ± 0.01 | -0.12 ± 0.01 | -0.86 ± 0.04 | -0.003 ± 0.02 |
| Amide sp<sup>2</sup>O | 3.4 ± 0.2 | 1.4 ± 0.1 | 0.75 ± 0.04 | 5.0 ± 0.3 | 0.70 ± 0.05 |
| Amide sp<sup>2</sup>N | -1.4 ± 0.1 | -0.50 ± 0.03 | -0.42 ± 0.02 | -2.0 ± 0.1 | -0.42 ± 0.03 |
| Amide sp<sup>2</sup>C | -3.1 ± 0.2 | -1.3 ± 0.07 | -0.77 ± 0.01 | -4.6 ± 0.3 | -0.36 ± 0.09 |

<sup>a</sup> cal<sup>-1</sup> mol<sup>-1</sup> molal<sup>-1</sup> Å<sup>-2</sup>

**Table S10.** Predicted Chemical Contributions to m-Values Quantifying Effects of NEP (PVP Model Monomer) and PVP Polymer Residues on Protein and α-Helix Unfolding.

| | Globular protein unfolding | α-helix unfolding |
| --- | --- | --- |
| NEP Monomer | -200 ± 40<sup>a</sup> | 1600 ± 100<sup>a</sup> |
| PVP Interior residue | -70 ± 20<sup>b</sup> | 690 ± 60<sup>b</sup> |
| PVP End residue | -270 ± 70<sup>b</sup> | 2400 ± 200<sup>b</sup> |

<sup>a</sup> cal mol<sup>-1</sup> molal<sup>-1</sup> for interaction with 1000 Å<sup>2</sup> of protein ΔASA.
<sup>b</sup> cal mol<sup>-1</sup> molal<sup>-1</sup> residue<sup>-1</sup> for 1000 Å<sup>2</sup> of protein ΔASA.

**Table S11.** Test of Dataset Size by Excluding µ_23_ Values for Selected Polar (urea, mad) and Nonpolar (1,3-deu, aama) Amides^a^.

| Amide Atom |  | Two-way Alpha Values (millical mol<sup>-1</sup> molal<sup>-1</sup> Å<sup>-4</sup>) |  |  |  |  |
| --- | --- | --- | --- | --- | --- | --- |
| i | j | Entire set of 105 $\mu_{23}$ | Exclude urea (79 $\mu_{23}$) | Exclude 1,3-deu (91 $\mu_{23}$) | Exclude aama (94 $\mu_{23}$) | Exclude 1,3-deu and mad (81 $\mu_{23}$) |
| sp<sup>2</sup>O | sp<sup>2</sup>C | -13.9 ± 3.8 | -13.4 ± 3.9 | -15.3 ± 5.4 | -14 ± 3.9 | -15.3 ± 3.3 |
| sp<sup>2</sup>O | sp<sup>2</sup>N | -10.8 ± 1.7 | -10.7 ± 2.1 | -12.5 ± 6.5 | -9.8 ± 2.3 | -11.4 ± 1.8 |
| sp<sup>2</sup>C | sp<sup>3</sup>C | -10.3 ± 0.5 | -10.5 ± 0.5 | -10.1 ± 0.6 | -10.3 ± 0.5 | -10.1 ± 0.4 |
| sp<sup>2</sup>C | sp<sup>2</sup>C | -11.1 ± 3.3 | -11.2 ± 3.4 | -10.3 ± 5 | -11.1 ± 3.4 | -10.3 ± 3.1 |
| sp<sup>3</sup>C | sp<sup>3</sup>C | -3.9 ± 0.1 | -3.9 ± 0.1 | -4 ± 0.4 | -3.9 ± 0.1 | -4.1 ± 0.1 |
| sp<sup>3</sup>C | sp<sup>2</sup>N | -3.8 ± 0.2 | -3.7 ± 0.2 | -3.6 ± 0.8 | -4.1 ± 0.2 | -3.8 ± 0.2 |
| sp<sup>2</sup>C | sp<sup>2</sup>N | 1.8 ± 1.3 | 2.5 ± 1.4 | 2.2 ± 1.8 | 1.8 ± 1.3 | 2.2 ± 1.2 |
| sp<sup>2</sup>N | sp<sup>2</sup>N | 3.4 ± 0.6 | 3.2 ± 0.9 | 4 ± 2.8 | 3 ± 0.9 | 3.8 ± 0.6 |
| sp<sup>3</sup>C | sp<sup>2</sup>O | 10.8 ± 0.5 | 11 ± 0.5 | 10.3 ± 2.2 | 11.6 ± 0.7 | 11 ± 0.6 |
| sp<sup>2</sup>O | sp<sup>2</sup>O | 18.1 ± 5.1 | 16.9 ± 5.4 | 23.2 ± 17.2 | 15.5 ± 6.7 | 19.8 ± 5.3 |

<sup>a</sup> Abbreviations as in Tables S3-4.

## Literature Cited

1. S. N. Timasheff, “Control of protein stability and reactions by weakly interacting cosolvents: The simplicity of the complicated” in Advances in Protein Chemistry, Vol 51: Linkage Thermodynamics of Macromolecular Interactions, E. DiCera, Ed. (Elsevier Academic Press Inc, San Diego, 1998), vol. 51, pp. 355–432.

2. M. T. Record, E. Guinn, L. Pegram, M. Capp, Introductory Lecture: Interpreting and predicting Hofmeister salt ion and solute effects on biopolymer and model processes using the solute partitioning model. Faraday Discuss. 160, 9–44 (2013).

3. K. A. Dill, Dominant forces in protein folding. Biochemistry 29, 7133–7155 (1990).

4. T. Kortemme, A. V. Morozov, D. Baker, An orientation-dependent hydrogen bonding potential improves prediction of specificity and structure for proteins and protein-protein complexes. J. Mol. Biol. 326, 1239–1259 (2003).

5. C. N. Pace, J. M. Scholtz, G. R. Grimsley, Forces stabilizing proteins. FEBS Lett. 588, 2177–2184 (2014).

6. R. W. Newberry, R. T. Raines, A prevalent intraresidue hydrogen bond stabilizes proteins. Nat. Chem. Biol. 12, 1084-+ (2016).

7. C. Tanford, The hydrophobic effect and the organization of living matter. Science 200, 1012–1018 (1978).

8. R. S. Spolar, M. T. Record, Jr., Coupling of local folding to site-specific binding of proteins to DNA. Science 263, 777–784 (1994).

9. R. L. Baldwin, Properties of hydrophobic free energy found by gas-liquid transfer. Proc. Natl. Acad. Sci. U. S. A. 110, 1670–1673 (2013).

10. R. L. Baldwin, Dynamic hydration shell restores Kauzmann’s 1959 explanation of how the hydrophobic factor drives protein folding. Proc. Natl. Acad. Sci. U. S. A. 111, 13052–13056 (2014).

11. J. Novotny, S. Bazzi, R. Marek, J. Kozelka, Lone-pair-pi interactions: analysis of the physical origin and biological implications. Phys. Chem. Chem. Phys. 18, 19472–19481 (2016).

12. G. J. Bartlett, A. Choudhary, R. T. Raines, D. N. Woolfson, n ->pi* interactions in proteins. Nat. Chem. Biol. 6, 615–620 (2010).

13. G. J. Bartlett, R. W. Newberry, B. VanVeller, R. T. Raines, D. N. Woolfson, Interplay of hydrogen bonds and n-->pi* interactions in proteins. J. Am. Chem. Soc. 135, 18682–18688 (2013).

14. R. W. Newberry, B. VanVeller, I. A. Guzei, R. T. Raines, n ->pi* Interactions of Amides and Thioamides: Implications for Protein Stability. Journal of the American Chemical Society 135, 7843–7846 (2013).

15. S. K. Singh, A. Das, The n -> pi* interaction: a rapidly emerging non-covalent interaction. Phys. Chem. Chem. Phys. 17, 9596–9612 (2015).

16. R. W. Newberry, R. T. Raines, The n ->pi* Interaction. Accounts Chem. Res. 50, 1838–1846 (2017).

17. R. W. Newberry, R. T. Raines, Secondary Forces in Protein Folding. ACS Chem. Biol. 14, 1677–1686 (2019).

18. E. A. Meyer, R. K. Castellano, F. Diederich, Interactions with aromatic rings in chemical and biological recognition. Angew. Chem.-Int. Edit. 42, 1210–1250 (2003).

19. A. J. Neel, M. J. Hilton, M. S. Sigman, F. D. Toste, Exploiting non-covalent pi interactions for catalyst design. Nature 543, 637–646 (2017).

20. E. S. Courtenay, M. W. Capp, C. F. Anderson, M. T. Record, Vapor pressure osmometry studies of osmolyte-protein interactions: Implications for the action of osmoprotectants in vivo and for the interpretation of “osmotic stress” experiments in vitro. Biochemistry 39, 4455–4471 (2000).

21. M. J. Plevin, D. L. Bryce, J. Boisbouvier, Direct detection of CH/pi interactions in proteins. Nat. Chem. 2, 466–471 (2010).

22. M. Nishio, Y. Umezawa, J. Fantini, M. S. Weiss, P. Chakrabarti, CH-pi hydrogen bonds in biological macromolecules. Phys. Chem. Chem. Phys. 16, 12648–12683 (2014).

23. K. L. Hudson et al., Carbohydrate-Aromatic Interactions in Proteins. Journal of the American Chemical Society 137, 15152–15160 (2015).

24. L. M. Pegram, M. T. Record, Thermodynamic origin of Hofmeister ion effects. J. Phys. Chem. B 112, 9428–9436 (2008).

25. M. W. Capp et al., Interactions of the Osmolyte Glycine Betaine with Molecular Surfaces in Water: Thermodynamics, Structural Interpretation, and Prediction of m-Values. Biochemistry 48, 10372–10379 (2009).

26. E. J. Guinn, L. M. Pegram, M. W. Capp, M. N. Pollock, M. T. Record, Quantifying why urea is a protein denaturant, whereas glycine betaine is a protein stabilizer. Proc. Natl. Acad. Sci. U. S. A. 108, 16932–16937 (2011).

27. R. C. Diehl, E. J. Guinn, M. W. Capp, O. V. Tsodikov, M. T. Record Jr, Quantifying additive interactions of the osmolyte proline with individual functional groups of proteins: comparisons with urea and glycine betaine, interpretation of m-values. Biochemistry 52, 5997–6010 (2013).

28. E. J. Guinn et al., Quantifying Functional Group Interactions That Determine Urea Effects on Nucleic Acid Helix Formation. Journal of the American Chemical Society 135, 5828–5838 (2013).

29. D. B. Knowles et al., Chemical Interactions of Polyethylene Glycols (PEGs) and Glycerol with Protein Functional Groups: Applications to Effects of PEG and Glycerol on Protein Processes. Biochemistry 54, 3528–3542 (2015).

30. X. Cheng et al., Basis of Protein Stabilization by K Glutamate: Unfavorable Interactions with Carbon, Oxygen Groups. Biophys. J. 111, 1854–1865 (2016).

31. X. Cheng et al., Experimental Atom-by-Atom Dissection of Amide-Amide and Amide-Hydrocarbon Interactions in H2O. Journal of the American Chemical Society 139, 9885–9894 (2017).

32. X. Cheng et al., Quantifying Interactions of Nucleobase Atoms with Model Compounds for the Peptide Backbone and Glutamine and Asparagine Side Chains in Water. Biochemistry 57, 2227–2237 (2018).

33. A. Ben-Naim, Inversion of Kirkwood-Buff theory of solutions - application to water-ethanol system. J. Chem. Phys. 67, 4884–4890 (1977).

34. R. Chitra, P. E. Smith, Preferential interactions of cosolvents with hydrophobic solutes. J. Phys. Chem. B 105, 11513–11522 (2001).

35. P. E. Smith, Chemical potential derivatives and preferential interaction parameters in biological systems from Kirkwood-Buff theory. Biophys. J. 91, 849–856 (2006).

36. D. R. Canchi, A. E. Garcia, “Cosolvent Effects on Protein Stability” in Annual Review of Physical Chemistry, Vol 64, M. A. Johnson, T. J. Martinez, Eds. (Annual Reviews, Palo Alto, 2013), vol. 64, pp. 273–293.

37. D. Trzesniak, N. F. A. van der Vegt, W. F. van Gunsteren, Computer simulation studies on the solvation of aliphatic hydrocarbons in 6.9 M aqueous urea solution. Phys. Chem. Chem. Phys. 6, 697–702 (2004).

38. D. Trzesniak, N. F. A. Van Der Vegt, W. F. Van Gunsteren, Analysis of neo-pentane-urea pair potentials of mean force in aqueous urea. Mol. Phys. 105, 33–39 (2007).

39. L. Ma, L. Pegram, M. T. Record, Q. Cui, Preferential Interactions between Small Solutes and the Protein Backbone: A Computational Analysis. Biochemistry 49, 1954–1962 (2010).

40. P. Ganguly, N. F. A. van der Vegt, Representability and Transferability of Kirkwood-Buff Iterative Boltzmann Inversion Models for Multicomponent Aqueous Systems. J. Chem. Theory Comput. 9, 5247–5256 (2013).

41. B. Lin, P. E. M. Lopes, B. Roux, A. D. MacKerell, Kirkwood-Buff analysis of aqueous N-methylacetamide and acetamide solutions modeled by the CHARMM additive and Drude polarizable force fields. J. Chem. Phys. 139, 11 (2013).

42. Y. Nozaki, C. Tanford, The Solubility of Amino Acids and Related Compounds in Aqueous Urea Solutions. J Biol Chem 238, 4074–4081 (1963).

43. M. Auton, D. W. Bolen, Additive transfer free energies of the peptide backbone unit that are independent of the model compound and the choice of concentration scale. Biochemistry 43, 1329–1342 (2004).

44. M. Auton, D. W. Bolen, Predicting the energetics of osmolyte-induced protein folding/unfolding. Proc. Natl. Acad. Sci. U. S. A. 102, 15065–15068 (2005).

45. M. Auton, L. M. F. Holthauzen, D. W. Bolen, Anatomy of energetic changes accompanying urea-induced protein denaturation. Proc. Natl. Acad. Sci. U. S. A. 104, 15317–15322 (2007).

46. M. Auton, J. Rosgen, M. Sinev, L. M. F. Holthauzen, D. W. Bolen, Osmolyte effects on protein stability and solubility: A balancing act between backbone and side-chains. Biophys. Chem. 159, 90–99 (2011).

47. E. J. Guinn, W. S. Kontur, O. V. Tsodikov, I. Shkel, M. T. Record, Jr., Probing the protein-folding mechanism using denaturant and temperature effects on rate constants. Proc Natl Acad Sci U S A 110, 16784–16789 (2013).

48. E. S. Courtenay, M. W. Capp, R. M. Saecker, M. T. Record, Thermodynamic analysis of interactions between denaturants and protein surface exposed on unfolding: Interpretation of urea and guanidinium chloride m-values and their correlation with changes in accessible surface area (ASA) using preferential interaction coefficients and the local-bulk domain model. Proteins, 72–85 (2000).

49. L. M. Pegram, M. T. Record, Partitioning of atmospherically relevant ions between bulk water and the water/vapor interface. Proc. Natl. Acad. Sci. U. S. A. 103, 14278–14281 (2006).

50. L. M. Pegram, M. T. Record, Hofmeister salt effects on surface tension arise from partitioning of anions and cations between bulk water and the air-water interface. J. Phys. Chem. B 111, 5411–5417 (2007).

51. L. M. Pegram, M. T. Record, Quantifying accumulation or exclusion of H(+), HO(-), and Hofmeister salt ions near interfaces. Chem. Phys. Lett. 467, 1–8 (2008).

52. L. M. Pegram, M. T. Record, Using Surface Tension Data to Predict Differences in Surface and Bulk Concentrations of Nonelectrolytes in Water. J. Phys. Chem. C 113, 2171–2174 (2009).

53. R. Deepak, R. Sankararamakrishnan, Unconventional N-H center dot center dot center dot N Hydrogen Bonds Involving Proline Backbone Nitrogen in Protein Structures. Biophys. J. 110, 1967–1979 (2016).

54. I. A. Shkel, D. B. Knowles, M. T. Record, Separating chemical and excluded volume interactions of polyethylene glycols with native proteins: Comparison with PEG effects on DNA helix formation. Biopolymers 103, 517–527 (2015).

55. D. B. Knowles, A. S. LaCroix, N. F. Deines, I. Shkel, M. T. Record, Separation of preferential interaction and excluded volume effects on DNA duplex and hairpin stability. Proc. Natl. Acad. Sci. U. S. A. 108, 12699–12704 (2011).

56. A. C. Miklos, C. G. Li, G. J. Pielak, “Using NMR-detected backbone amide H-1 exchange to assess macromolecular crowding effects on globular-protein stability” in Methods in Enzymology, Vol 466: Biothermodynamics, Pt B, M. L. Johnson, J. M. Holt, G. K. Ackers, Eds. (Elsevier Academic Press Inc, San Diego, 2009), vol. 466, pp. 1-+.

57. L. M. Charlton et al., Residue-level interrogation of macromolecular crowding effects on protein stability. Journal of the American Chemical Society 130, 6826–6830 (2008).

58. A. C. Miklos, C. G. Li, N. G. Sharaf, G. J. Pielak, Volume Exclusion and Soft Interaction Effects on Protein Stability under Crowded Conditions. Biochemistry 49, 6984–6991 (2010).

59. R. J. Hefford, Polymer Mixing in Aqueous-Solution. Polymer 25, 979–984 (1984).

60. K. A. Dill, S. Bromberg, D. Stigter, Molecular driving forces: statistical thermodynamics in biology, chemistry, physics, and nanoscience 2 (Garland Science, 2012).

61. J.-M. Choi, A. S. Holehouse, R. V. Pappu, Physical Principles Underlying the Complex Biology of Intracellular Phase Transitions. Annual Review of Biophysics 49, 107–133 (2020).

62. M. Feric et al., Coexisting Liquid Phases Underlie Nucleolar Subcompartments. Cell 165, 1686–1697 (2016).

63. H. Hofmann et al., Polymer scaling laws of unfolded and intrinsically disordered proteins quantified with single-molecule spectroscopy. Proc. Natl. Acad. Sci. U. S. A. 109, 16155–16160 (2012).

64. A. Soranno et al., Single-molecule spectroscopy reveals polymer effects of disordered proteins in crowded environments. Proc. Natl. Acad. Sci. U. S. A. 111, 4874–4879 (2014).

65. J. A. Riback et al., Stress-Triggered Phase Separation Is an Adaptive, Evolutionarily Tuned Response. Cell 168, 1028-+ (2017).

66. Y. Shin, C. P. Brangwynne, Liquid phase condensation in cell physiology and disease. Science 357, 7 (2017).

67. T. M. Franzmann et al., Phase separation of a yeast prion protein promotes cellular fitness. Science 359, 47-+ (2018).

## References

1. J. Sadowski, J. Gasteiger, G. Klebe, Comparison of Automatic 3-Dimensional Model Builders Using 639 X-Ray Structures. J Chem Inf Comp Sci 34, 1000–1008 (1994).

2. O. V. Tsodikov, M. T. Record, Jr., Y. V. Sergeev, Novel computer program for fast exact calculation of accessible and molecular surface areas and average surface curvature. Journal of computational chemistry 23, 600–609 (2002).

3. J. R. Livingstone, R. S. Spolar, M. T. Record, Jr., Contribution to the thermodynamics of protein folding from the reduction in water-accessible nonpolar surface area. Biochemistry 30, 4237–4244 (1991).

4. E. E. Bolton et al., PubChem3D: a new resource for scientists. J Cheminformatics 3 (2011).

5. E. L. Ulrich et al., BioMagResBank. Nucleic Acids Res 36, D402–D408 (2008).

6. W. Humphrey, A. Dalke, K. Schulten, VMD: visual molecular dynamics. J Mol Graph 14, 33-38, 27-38 (1996).

7. R. Fraczkiewicz, W. Braun, Exact and efficient analytical calculation of the accessible surface areas and their gradients for macromolecules. Journal of computational chemistry 19, 319–333 (1998).

8. X. Cheng et al., Quantifying Interactions of Nucleobase Atoms with Model Compounds for the Peptide Backbone and Glutamine and Asparagine Side Chains in Water. Biochemistry 57, 2227–2237 (2018).

9. X. Cheng et al., Experimental Atom-by-Atom Dissection of Amide-Amide and Amide-Hydrocarbon Interactions in H2O. Journal of the American Chemical Society 139, 9885–9894 (2017).

10. I. A. Shkel, D. B. Knowles, M. T. Record, Separating chemical and excluded volume interactions of polyethylene glycols with native proteins: Comparison with PEG effects on DNA helix formation. Biopolymers 103, 517–527 (2015).

11. E. J. Guinn, L. M. Pegram, M. W. Capp, M. N. Pollock, M. T. Record, Jr., Quantifying why urea is a protein denaturant, whereas glycine betaine is a protein stabilizer. Proceedings of the National Academy of Sciences of the United States of America 108, 16932–16937 (2011).

12. E. J. Guinn, L. M. Pegram, M. W. Capp, M. N. Pollock, M. T. Record, Quantifying why urea is a protein denaturant, whereas glycine betaine is a protein stabilizer. Proc. Natl. Acad. Sci. U. S. A. 108, 16932–16937 (2011).

13. D. B. Knowles et al., Chemical Interactions of Polyethylene Glycols (PEGs) and Glycerol with Protein Functional Groups: Applications to Effects of PEG and Glycerol on Protein Processes Biochemistry 54, 3528–3542 (2015).

